# Genome-wide arrayed CRISPR activation screen for prion protein modulators

**DOI:** 10.64898/2026.03.01.707423

**Authors:** Chiara Trevisan, Hao Wang, Vangelis Bouris, Simon Mead, Jiang-An Yin, Adriano Aguzzi

**Affiliations:** Institute of Neuropathology, University of Zurich, CH-8091 Zurich, Switzerland; Institute for the Science of the Aging Brain (ISAB), CH-9004 St. Gallen, Switzerland; Institute of Prion Diseases, MRC Prion Unit at University College London, London, UK; Present address: Aging and Regeneration Center, School of Basic Medical Sciences, Tsinghua University, 100084, Beijing, P.R. China

**Author notes:** Equal contribution (Chiara Trevisan, Hao Wang).

## Abstract

The cellular prion protein (PrP^C^) is an essential substrate for prion propagation, and its abundance strongly influences susceptibility to prion disease. To systematically identify genetic regulators of PrP^C^ abundance, we performed an arrayed genome-wide CRISPR activation (CRISPRa) screen targeting 19,839 human protein-coding genes in human glioblastoma cells. Using a quantitative time-resolved fluorescence resonance energy transfer (TR-FRET) immunoassay, we measured PrP^C^ levels across 22,442 individual genetic perturbations. This screen identified 531 genes whose activation significantly modulated PrP^C^ abundance. A curated subset of 50 candidates was subjected to validation using independent TR-FRET assays and Western blotting, confirming 45 (90%) of tested hits. All raw and processed screening data, metadata, and analysis code are publicly available. This dataset provides a comprehensive resource for studying genetic regulation of PrP^C^ homeostasis and enables reuse for investigations of membrane protein trafficking, proteostasis, and prion biology.

## Background and summary

Prion diseases are rare but invariably fatal neurodegenerative disorders affecting humans and animals, including Creutzfeldt–Jakob disease (CJD), fatal familial insomnia, kuru, and scrapie^1,2^. These disorders are characterized by progressive neuronal loss, spongiform degeneration, and gliosis, leading to rapid cognitive decline, motor dysfunction, and death, often within months of clinical onset^1,3,4^. At present, there are no disease-modifying therapies for prion diseases, and treatment remains purely supportive. Because prion propagation requires the cellular prion protein (PrP^C^) as an obligate substrate, PrP^C^ abundance is a key rate-limiting factor in disease progression^5^. Genetic ablation or reduction of PrP^C^ is strongly protective in multiple experimental models^6,7,7–13^, establishing PrP^C^ dosage as a central node in prion biology and a compelling therapeuric target.

Despite the importance of PrP^C^ abundance, the cellular and genetic mechanisms that govern PrP^C^ homeostasis remain incompletely understood. PrP^C^ levels reflect the net outcome of multiple regulatory processes, including gene expression, protein maturation, membrane localization, and clearance, which are likely coordinated by diverse and context-dependent regulatory programs.

However, these programs have not been systematically mapped at genome scale, limiting our ability to understand how PrP^C^ abundance is dynamically regulated across cellular states.

To address this gap, we performed an arrayed genome-wide CRISPR activation (CRISPRa) screen in human glioblastoma cells (Fig. 1A) using the T.gonfio quadruple-guide RNA library (Fig. 1B), targeting all annotated human protein-coding genes. We selected the human glioblastoma cell line U-251 MG because it exhibits intermediate endogenous PrP^C^ expression levels (Fig. 2A,B), enabling the detection of both positive and negative genetic regulators of PrP^C^ abundance. Stable dCas9-VPR expression and PRNP activation controls were used to benchmark CRISPRa performance and define the readout window and screening conditions (Fig. 2C,D).

**Figure 1.**
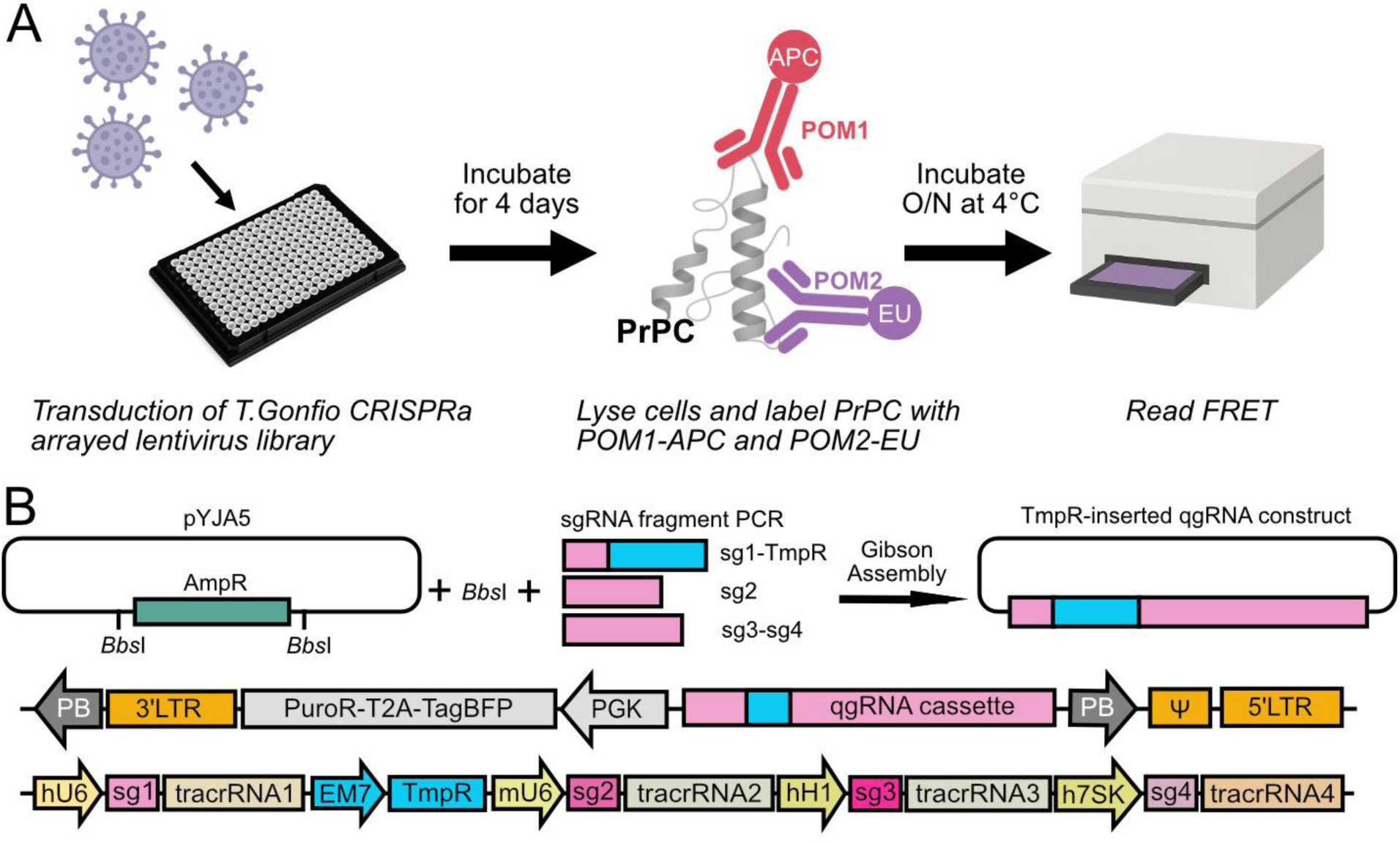
Overview of the arrayed genome-wide CRISPRa screen and qgRNA library design. (A) Schematic of the arrayed CRISPR activation (CRISPRa) screen performed in U-251 MG cells stably expressing dCas9-VPR. Cells were transduced with the T.gonfio quadruple-guide RNA (qgRNA) lentiviral library, targeting human protein-coding genes at single-gene resolution. PrP^C^ abundance was quantified four days post-transduction using a solution-based time-resolved fluorescence resonance energy transfer (TR-FRET) immunoassay. (B) Schematic of the qgRNA-pYJA5 construct and cloning strategy underlying the T.gonfio CRISPRa library (adapted from Yin et al., Nat. Biomed. Eng., 2025^14^). The ampicillin resistance gene (AmpR) was removed from the parental pYJA5 vector. sgRNA1–4 and the trimethoprim resistance gene (TmpR) were generated as three distinct PCR amplicons and assembled by Gibson cloning to generate the qgRNA-pYJA5 plasmid. Transformants were selected using trimethoprim. The full plasmid structure and detailed organization of the qgRNA cassette are shown. LTR, long terminal repeat; Ψ, packaging signal; PB, piggyBac transposon element; PuroR, puromycin resistance gene; hU6, mU6, hH1, and h7SK, RNA polymerase III promoters; sg, single-guide RNA. F and R arrows indicate primer positions used for single-colony PCR, Sanger sequencing, and next-generation sequencing validation.

**Figure 2.**
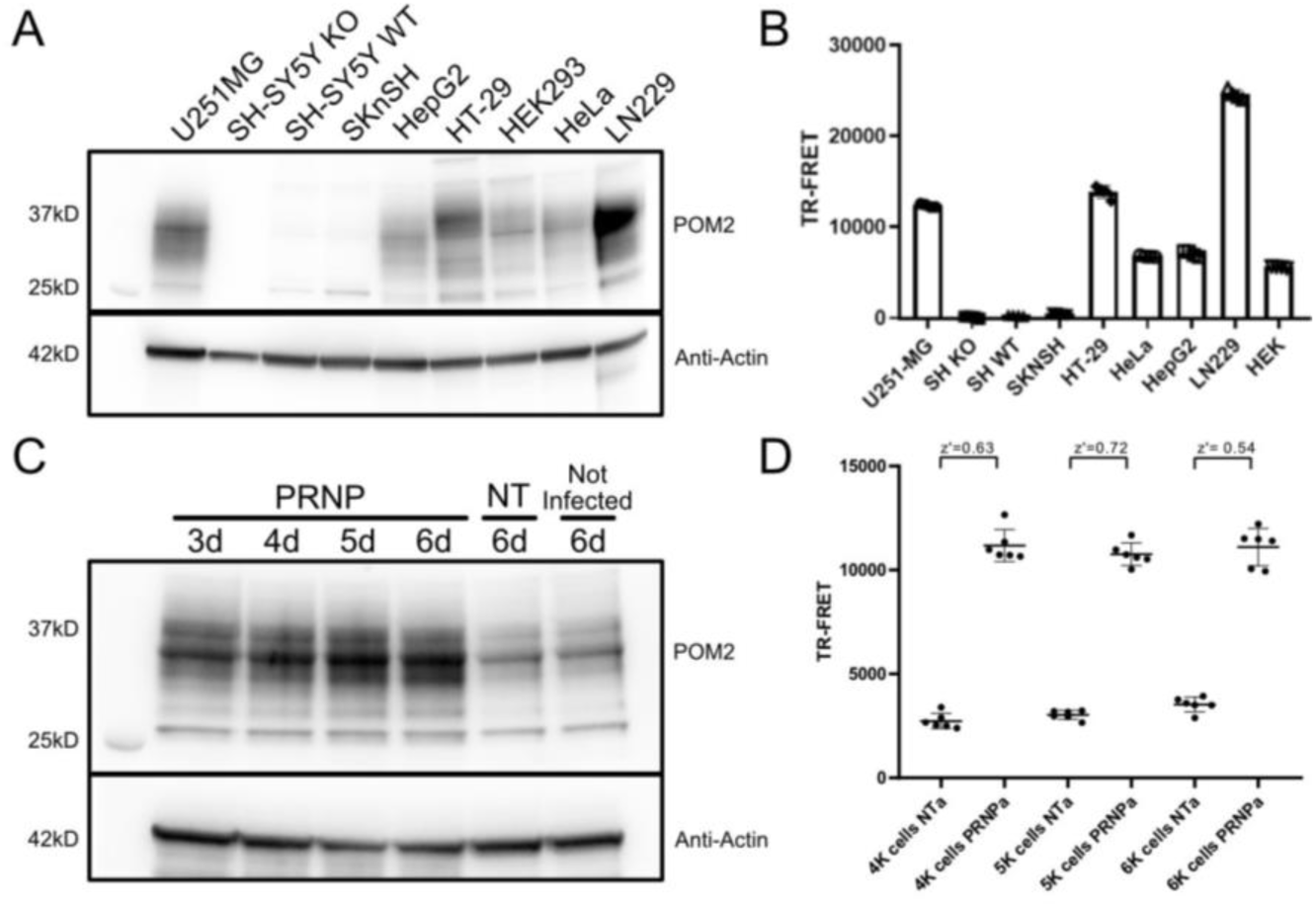
Establishment and optimization of the CRISPRa screening platform for PrP^C^ quantification. (A) Western blot analysis of PrP^C^ expression in lysates from the indicated human cell lines using the POM2 antibody. The cell lines tested include U-251 MG, SH-SY5Y wild type (SH WT), SH-SY5Y PRNP knockout (SH KO), SK-N-SH, LN229, HEK293, HeLa, HepG2, and HT-29. Actin was used as a loading control. (B) TR-FRET–based quantification of PrP^C^ levels in the same panel of cell lines shown in (A). Data represent mean ± SEM from four independent measurements. U-251 MG cells exhibit intermediate PrP^C^ expression, enabling detection of both positive and negative regulators in CRISPRa screens. (C) Western blot analysis demonstrating CRISPRa-mediated overexpression of PrP^C^ in U-251 MG dCas9-VPR cells transduced with qgRNAs targeting PRNP or non-targeting (NT) controls. PrP^C^ was detected using the POM2 antibody, and actin served as a loading control. PrP^C^ induction was monitored over time post-transduction. (D) TR-FRET–based assay optimization testing different cell seeding densities at a multiplicity of infection (MOI) of 3. qgRNAs targeting PRNP served as positive controls and NT qgRNAs as negative controls. Z′-factor analysis was used to determine optimal screening conditions. Data are shown as sextuplicate measurements.

We employed a CRISPRa-based gain-of-function strategy rather than a loss-of-function approach to comprehensively interrogate regulatory mechanisms controlling PrP^C^ levels. CRISPRa enables the activation of genes that are transcriptionally silent or expressed at low levels under basal conditions, including regulatory factors whose functions may not be revealed by knockout strategies. In addition, gain-of-function perturbations can uncover regulatory logic that is buffered by redundancy in loss-of-function screens and allow direct testing of whether increased gene activity is sufficient to modulate PrP^C^ abundance. This approach therefore provides complementary and mechanistically informative insight into PrP^C^ regulation.

A central strength of this approach is the use of T.gonfio, a novel arrayed CRISPRa library^14^(Fig. 1B) providing precise, well-controlled gain-of-function perturbations for over 20,000 genes. Each gene was targeted using quadruple single-guide RNAs per transcription start site, designed to tolerate common human polymorphisms. This design maximizes on-target activation, minimizes dropout due to sequence variability, and enables robust induction across a wide spectrum of gene expression states.

The screen was implemented in 384-well plates with dedicated layout features to control for positional effects and assay-specific background. Control placement and TR-FRET antibody calibration wells are summarized in Fig. 3A-B, and a representative plate-level heatmap illustrates the spatial distribution of the net TR-FRET signal and internal controls used for quality metrics (Fig. 3C).

**Figure 3.**
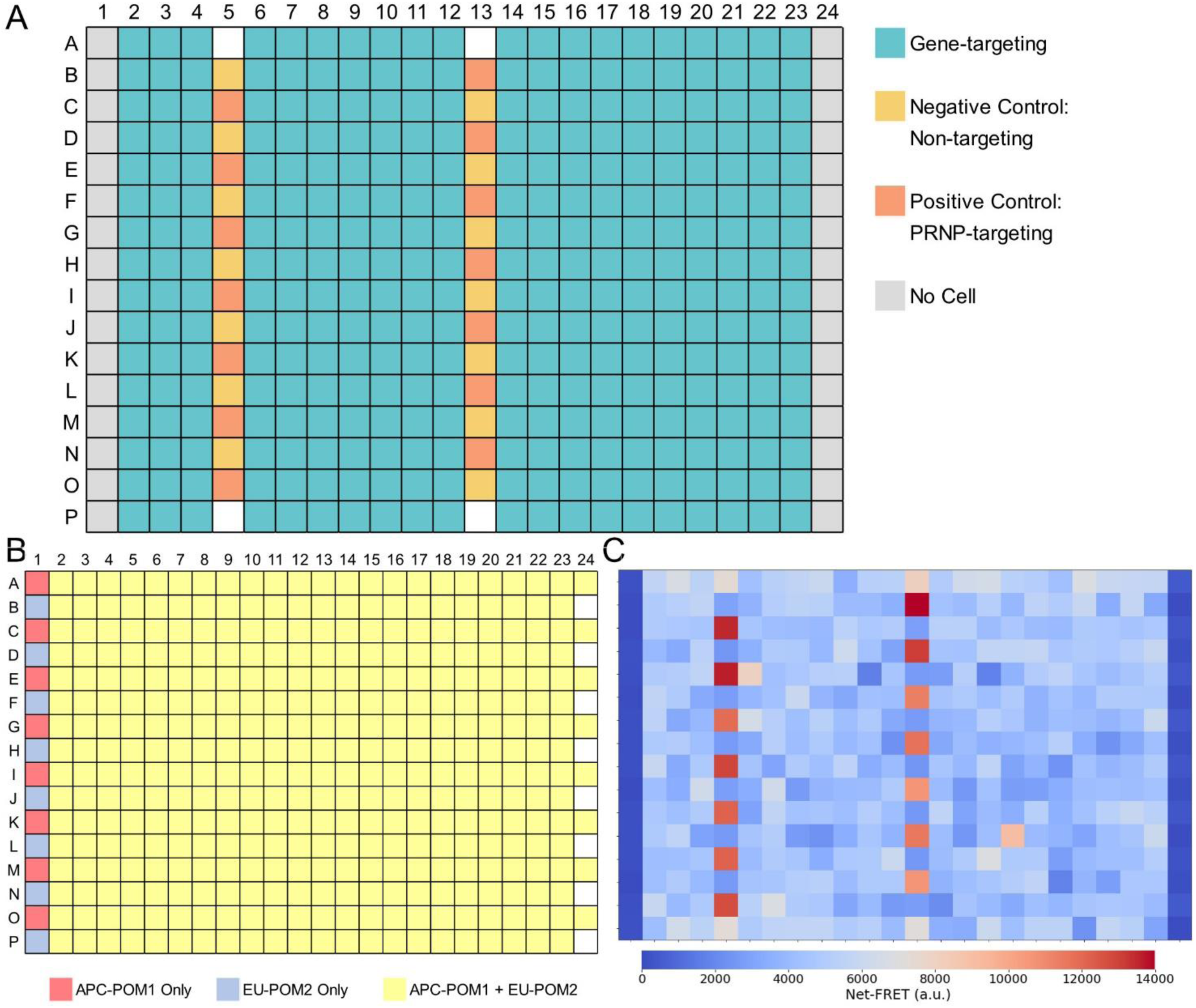
384-well plate layouts for genetic perturbations and TR-FRET assay configuration. **A.** Schematic of the 384-well plate layout used for arrayed CRISPRa genetic perturbations. Columns 1 and 24 contain medium only and no cells and were reserved for TR-FRET normalization. Columns 5 and 13 were designated as control columns and contained alternating non-targeting controls (NTC) and PRNP-targeting positive controls arranged to control for positional effects. In column 5, wells B, D, F, H, J, L, and N contained NTCs, while wells C, E, G, I, K, M, and O contained PRNP-targeting controls; in column 13, this pattern was inverted, with PRNP-targeting controls in wells B, D, F, H, J, L, and N and NTCs in wells C, E, G, I, K, M, and O. Placement of control columns at two distinct positions within the plate enabled monitoring and correction of potential spatial effects on TR-FRET signal acquisition. (B) Schematic of the 384-well plate layout used for TR-FRET antibody addition. Columns 1 and 24 were reserved for assay normalization and background control. In column 1, wells A, C, E, G, I, K, M, and O contained APC-POM1 only, while wells B, D, F, H, J, L, N, and P contained Eu-POM2 only. In column 24, wells A, C, E, G, I, K, M, and O contained both APC-POM1 and Eu-POM2, while wells B, D, F, H, J, L, N, and P contained no antibodies. These configurations enabled independent measurement of donor-only and acceptor-only background fluorescence, assessment of bleed-through and cross-excitation, and accurate calculation of net TR-FRET signal. All remaining wells contained both APC-POM1 and Eu-POM2 to quantify PrP^C^ abundance. (C) Representative heatmap of a single 384-well plate from the genome-wide CRISPRa screen, showing raw net TR-FRET signal (Net-FRET, arbitrary units) for all wells arranged according to the layouts in (A) and (B). Color intensity reflects absolute net TR-FRET signal, with warmer colors indicating higher PrPᶜ abundance and cooler colors indicating lower signal. Assay robustness for this plate, calculated using PRNP-targeting positive controls and non-targeting controls in columns 5 and 13, yielded a Z′ factor of 0.62, indicating excellent assay performance suitable for high-throughput screening.

PrP^C^ abundance was quantified using a sensitive TR-FRET immunoassay^15^ across 44,884 measurements corresponding to 22,442 individual genetic perturbations assayed in duplicate, resulting in the identification of 531 candidate genes that significantly modulate PrP^C^ levels (Fig. 4). Assay performance and reproducibility were assessed using plate-wise quality metrics and replicate concordance (Fig. 5), supporting the robustness of the dataset. A curated subset of 50 candidates, based on the strength of their effects and relevance to neuronal expression profiles was further validated (Fig. 6). Of these, 45 out of 50 candidates (90%) were independently validated, demonstrating a high confirmation rate and underscoring the robustness and reproducibility of the primary screen.

**Figure 4.**
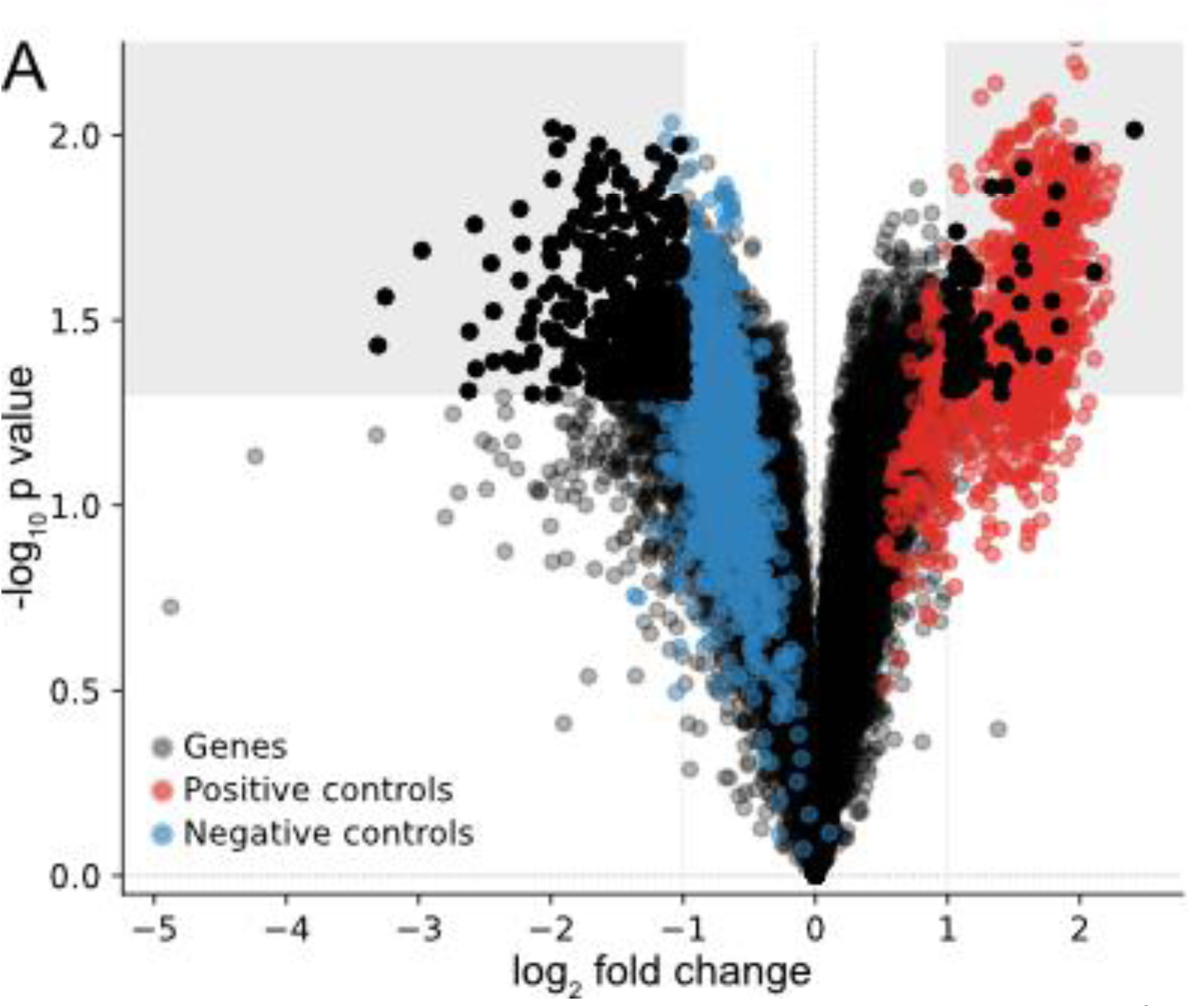
Genome-wide CRISPRa screen identifies genetic modulators of PrP^C^ abundance. (A) Volcano plot displaying log₂ fold change (log₂FC) versus –log₁₀(P value) for all genes tested in the arrayed CRISPRa screen. Each point represents a single genetic perturbation. Non-targeting (NT) control qgRNAs are highlighted in blue, and PRNP-targeting qgRNAs are highlighted in red.

**Figure 5.**
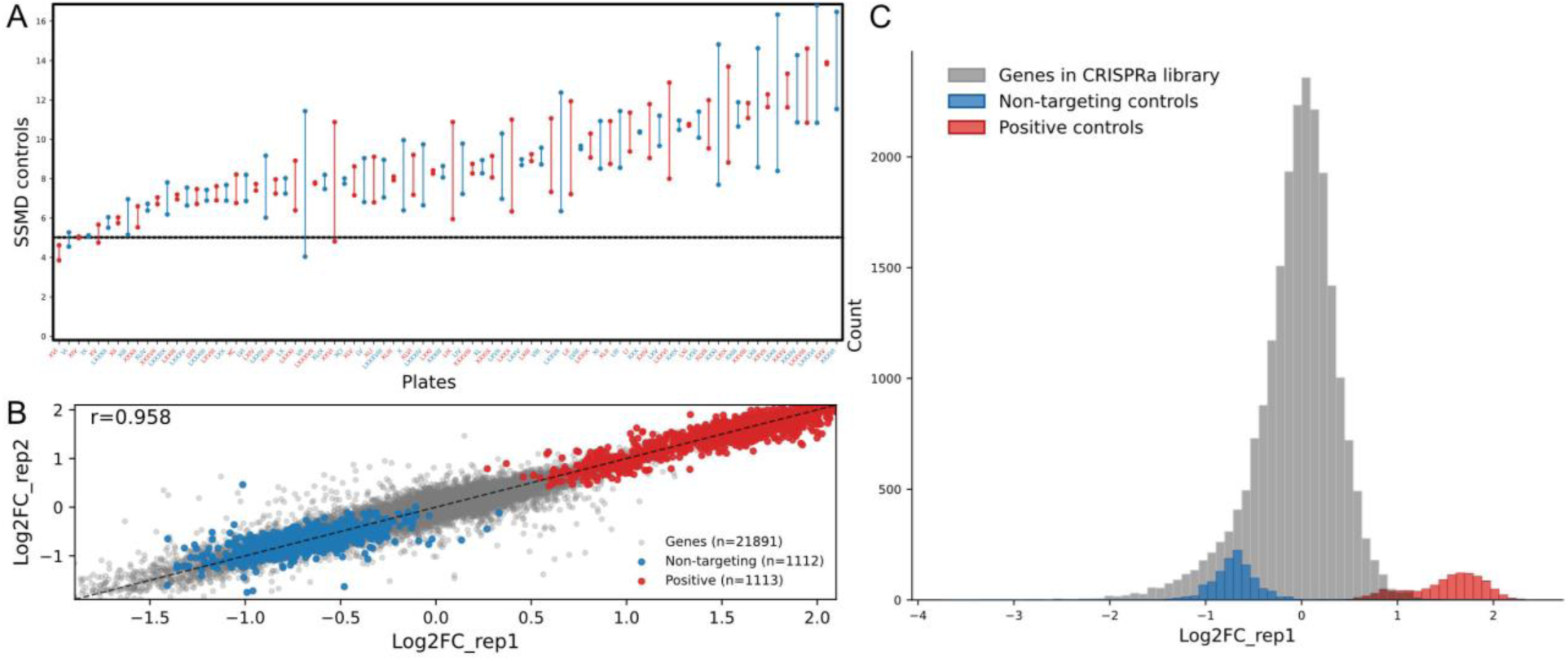
Assay performance and reproducibility of the genome-wide CRISPRa screen. (A) Plate-level assay quality assessed by strictly standardized mean difference (SSMD), calculated from PRNP-targeting positive controls and non-targeting (NT) controls on each 384-well plate. SSMD values quantify effect separation while accounting for variance and are well suited for duplicate-measurement screen designs. (B) Scatter plot showing strong reproducibility of replicate measurements, with log₂ fold-change (log₂FC) values from replicate 1 plotted against replicate 2 for all gene-targeting perturbations. (C) Histogram illustrating the distribution of log2FC fluorescence values across three groups: NT controls, PRNP-targeting controls and screened genes.

**Figure 6.**
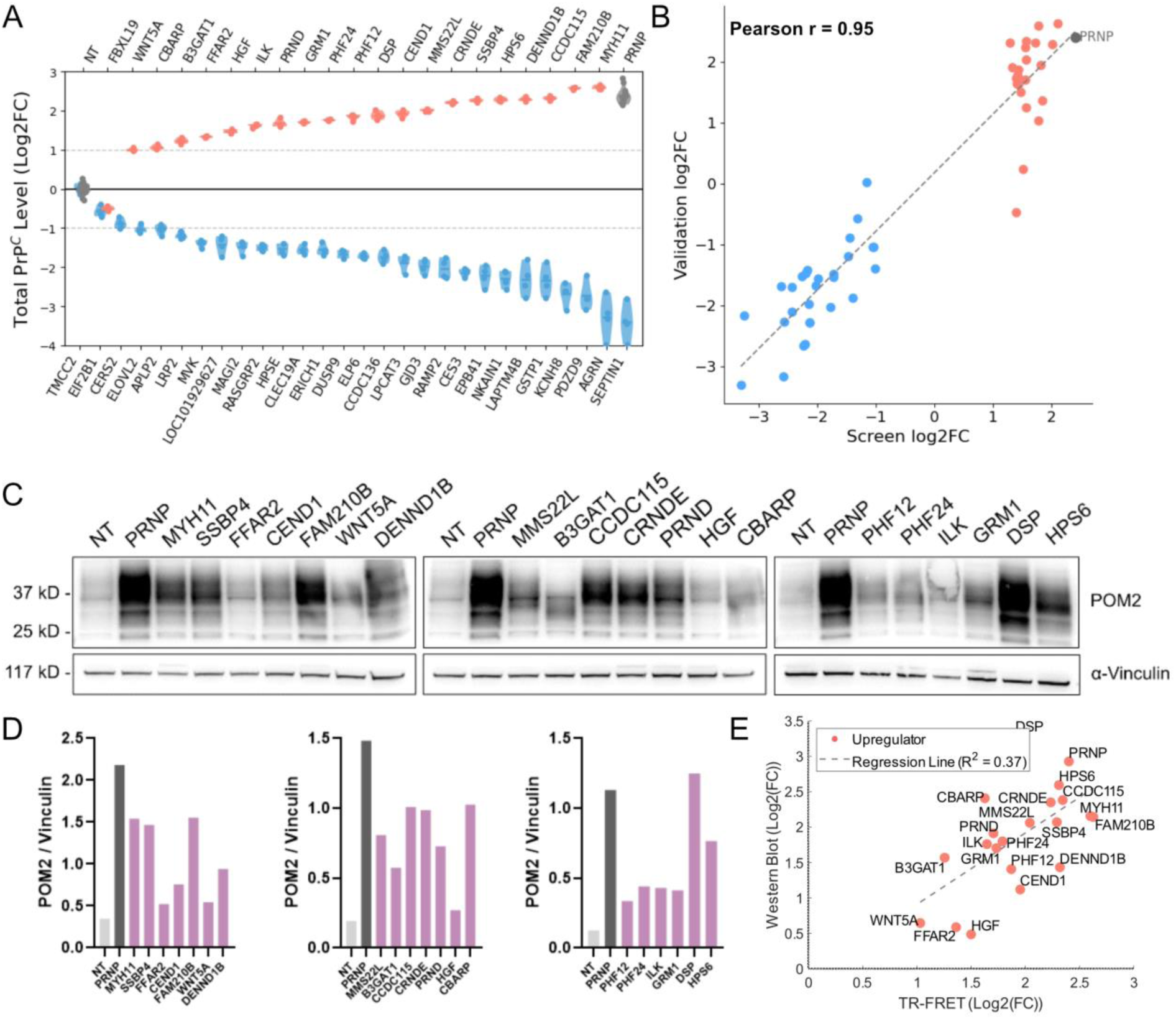
Secondary validation and cross-assay concordance of CRISPRa-identified PrP^C^ modulators. (A) Secondary TR-FRET–based quantification of PrP^C^ abundance for 50 candidate genes identified in the primary genome-wide CRISPRa screen. Each condition was measured with ≥8 technical replicates. Non-targeting (NT) and PRNP-targeting positive controls are shown in grey; PrP^C^ upregulators are shown in red; PrP^C^ downregulators are shown in blue. (B) Correlation between primary screen effect sizes and secondary TR-FRET validation results for the same 50 candidate genes. The x-axis shows log₂ fold-change (log₂FC) values from the primary screen, and the y-axis shows log₂FC values from secondary TR-FRET validation. Pearson correlation coefficient r = 0.95. PRNP-targeting controls are shown in grey and labeled; upregulators are shown in red; downregulators are shown in blue. (C) Immunoblot analysis of selected PrP^C^ upregulators following CRISPRa activation. PrP^C^ was detected using the POM2 antibody; vinculin served as a loading control. (D) Quantification of Western blot–derived PrP^C^ protein levels normalized to vinculin and expressed relative to non-targeting (NT) controls. (E) Correlation between TR-FRET measurements and Western blot quantification of total PrP^C^ levels for validated upregulators. The x-axis represents TR-FRET log₂FC values and the y-axis represents normalized Western blot log₂FC values.

Collectively, this dataset defines a comprehensive genetic landscape of PrP^C^ regulation and provides a framework for dissecting the molecular logic that controls PrP^C^ homeostasis. By capturing regulators that influence PrP^C^ abundance through diverse mechanisms, including protein production, stability, and turnover, this study offers an unbiased resource for understanding how PrP^C^ levels are set and maintained in human cells. More broadly, the gain-of-function nature of the screen enables exploration of regulatory programs that may be engaged under dynamic cellular conditions, such as changes in signaling state, tissue context, or environmental cues, providing insight into how PrP^C^ regulation may be modulated in physiological and disease-relevant settings.

Key assay parameters, control configuration, and screening outcomes are summarized in Table 1.

**Table 1.**
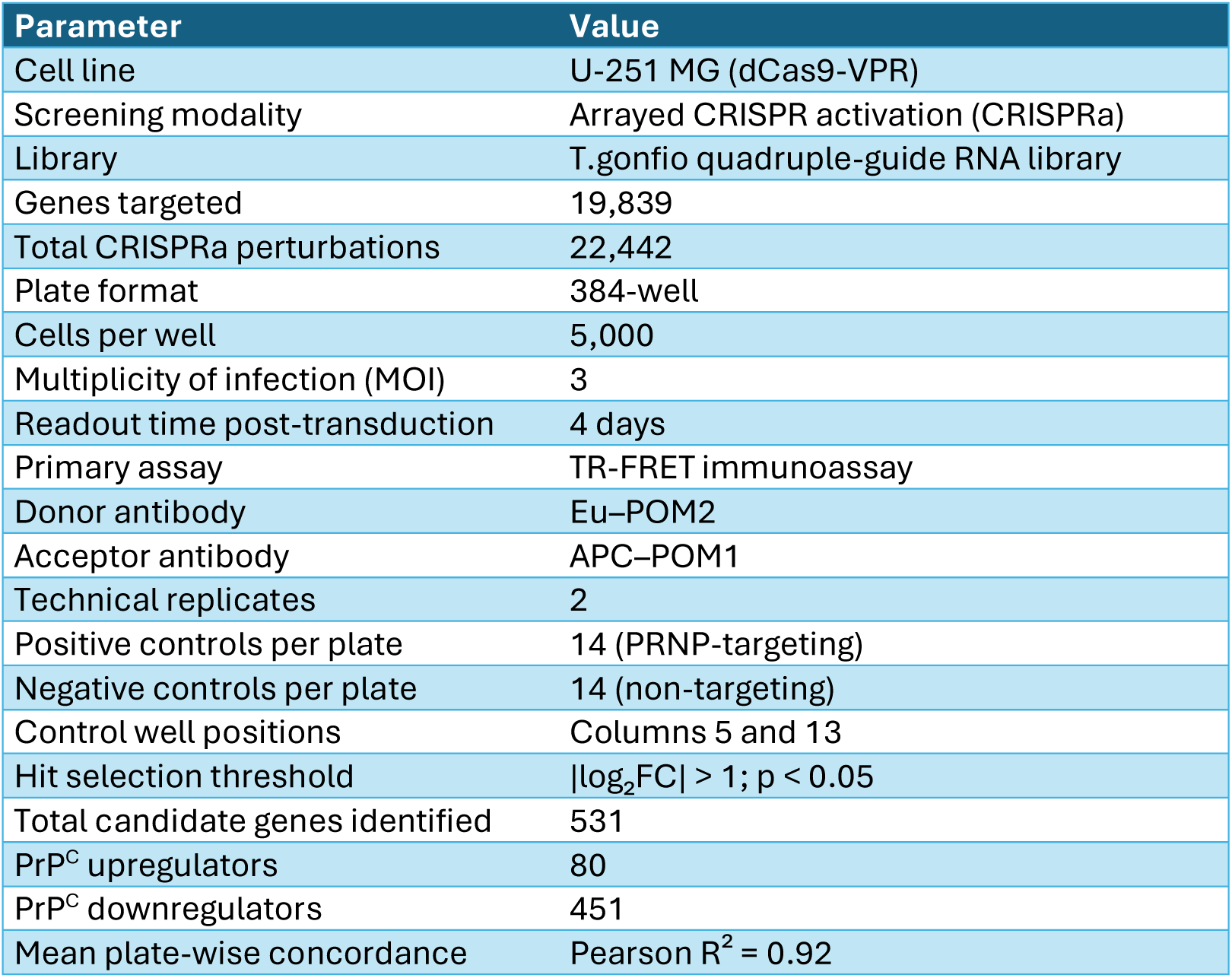
Design, performance, and outcomes of the genome-wide CRISPRa screen for PrP^C^ regulators. Summary of key experimental parameters, assay configuration, and screening outcomes for the arrayed genome-wide CRISPR activation (CRISPRa) screen performed in U-251 MG cells stably expressing dCas9-VPR. The screen targeted 19,839 human protein-coding genes using the T.gonfio quadruple-guide RNA library and quantified PrP^C^ abundance four days post-transduction by a TR-FRET–based immunoassay employing Eu–POM2 and APC–POM1 antibodies. Each perturbation was assayed in duplicate in 384-well plates containing PRNP-targeting positive controls and non-targeting negative controls positioned in columns 5 and 13. Candidate genes were defined based on an absolute log₂ fold-change (log₂FC) greater than 1 and a p-value < 0.05. Plate-wise reproducibility was assessed by Pearson correlation analysis.

## Methods

### Cell culture

Human glioblastoma U-251 MG cells (Kerafast, Inc., Boston, MA, USA; Accession ID: CVCL_0021) stably expressing dCas9-VPR (pXPR_120, Addgene #96917) were maintained in T150 tissue culture flasks (TPP, Trasadingen, Switzerland). Cells were cultured in OptiMEM without phenol red (Gibco, Thermo Fisher Scientific, Waltham, MA, USA) supplemented with 10% fetal bovine serum (FBS; Takara, Göteborg, Sweden), 1% non-essential amino acids (NEAA; Gibco), 1% GlutaMAX (Gibco), 1% penicillin–streptomycin (P/S; Gibco), and 10 µg/mL blasticidin (Gibco).

For genome-wide screening, polyclonal U-251 MG cells stably expressing endonuclease-deficient Cas9 (dCas9) fused to the tripartite transcriptional activator VP64-p65-RTA (VPR) were expanded under standard culture conditions, washed with phosphate-buffered saline (PBS; Gibco), and dissociated using Accutase (Gibco). Cells were resuspended in OptiMEM without phenol red, pooled, and counted using a TC20 Automated Cell Counter (Bio-Rad) with trypan blue staining (Gibco) to assess viability. Phenol red was excluded from the culture medium because it exhibits fluorescence and absorbance in spectral ranges that can interfere with time-resolved fluorescence resonance energy transfer (TR-FRET) measurements, potentially increasing background signal and reducing assay sensitivity and dynamic range. The use of phenol-red–free medium therefore improves the accuracy and reproducibility of TR-FRET–based quantification of PrPC abundance. All cells used in the primary screen were maintained at the same passage number to ensure experimental consistency. For long-term storage, cells were cryopreserved in Bambanker™ freezing medium (GC Lymphotec Inc.).

U-251 MG cells were selected for screening because they exhibit intermediate endogenous PrP^C^ expression relative to other commonly used human cell lines, enabling sensitive detection of both PrP^C^ upregulation and downregulation in CRISPRa-based perturbation screens (Fig. 2A,B).

### T.gonfio lentiviral library

The T.gonfio library is a genome-wide, arrayed CRISPR activation (CRISPRa) resource designed to enable robust and reproducible transcriptional upregulation of human protein-coding genes at single-gene resolution. The library is available in both plasmid-based and lentiviral formats. The plasmid library is distributed in 384-well PCR plates, with each well containing DNA at a concentration of 30 ng/µL, while the lentiviral version is organized across 65 plates of the same format. To ensure consistency and facilitate quality control in arrayed screening workflows, columns 5 and 13 of each plate are reserved for control wells and contain no genetic constructs. A schematic of the qgRNA vector architecture and cloning strategy is provided for reference (Fig. 1B) to facilitate reproducibility and reuse of the dataset.

Each T.gonfio construct encodes a quadruple-guide RNA (qgRNA) cassette comprising four distinct single-guide RNAs (sgRNAs) targeting the same gene or transcription start site (TSS). The four sgRNAs are expressed under the control of four different RNA polymerase III promoters (human U6, mouse U6, human H1, and human 7SK), a configuration that enhances activation potency and reduces variability across cell types and experimental conditions.

The library was generated using an automated liquid-phase assembly (ALPA) cloning strategy, enabling high-throughput, colony-free construction of arrayed qgRNA plasmids. sgRNAs were selected using a custom computational pipeline that integrates predicted on-target activity, GuideScan specificity scores, avoidance of common human genetic polymorphisms, and enforced spatial separation between guides to promote synergistic CRISPRa-mediated transcriptional activation. For genes with multiple annotated transcription start sites, independent qgRNA constructs were designed to target individual TSSs, allowing precise activation of alternative promoters.

In total, the T.gonfio library comprises 22,442 plasmids targeting 19,839 human protein-coding genes. The distribution of plasmids across functional sublibraries and overall library composition are summarized in Table 2. Each construct is embedded in a lentiviral-compatible backbone containing selection and fluorescent reporter elements (Fig. 1B), enabling efficient delivery by transfection or viral transduction and supporting standardized arrayed screening workflows.

**Table 2.** Composition of the T.gonfio genome-wide arrayed CRISPRa library by sublibrary. Counts of quadruple-guide RNA (qgRNA) plasmids distributed across functional sublibraries within the T.gonfio resource. The table reports the number of plasmid constructs per sublibrary, together with the total number of protein-coding genes targeted, the total number of targeted transcription start sites/open reading frames (TSS/ORF), the number of non-targeting qgRNA control plasmids, and the overall number of plasmids in the full library. Because some genes are represented by multiple promoter annotations, the number of targeted TSS/ORF exceeds the number of genes.

| Sublibrary | #Plasmids |
| --- | --- |
| Transcription factors | 1634 |
| GPCRs | 802 |
| Secretome | 1694 |
| Membrane Proteins (non-GPCR) | 1760 |
| Kinases/Phosphatases/Drug Targets | 2012 |
| Mitochondria/Trafficking/Motility | 2052 |
| Stress/Proteostasis | 2699 |
| Cancer/Apoptosis | 2266 |
| Gene Expression (non-TF) | 1131 |
| Unassigned | 3789 |
| **Library Summary** |  |
| Total genes targeted | 19839 |
| Total TSS/ORF targeted | 22326 |
| Non-targeting 4sgRNA plasmid controls | 116 |
| **Total plasmids in the library** | **22442** |

The quadruple-guide architecture of T.gonfio enables reliable activation of both highly expressed and transcriptionally low or silent genes, a key advantage for gain-of-function screens aimed at uncovering regulators that are inactive under basal conditions. As such, the T.gonfio CRISPRa library provides a scalable and versatile platform for genome-wide activation screens, particularly for phenotypes that are not accessible using pooled loss-of-function approaches. For a comprehensive overview of the T.gonfio library’s design and applications, refer to its original publication^14^.

### Lentiviral packaging

HEK293T cells were maintained in Dulbecco’s Modified Eagle Medium (DMEM; Gibco, Thermo Fisher Scientific, Waltham, MA, USA) supplemented with 10% fetal bovine serum (FBS) and cultured in poly-D-lysine–coated six-well plates until reaching 80–90% confluency. Lentiviral particles were generated by transient transfection using Lipofectamine™ 3000 (Invitrogen, Thermo Fisher Scientific, Carlsbad, CA, USA) with the CRISPRa transfer plasmid together with the packaging plasmid pAX-2 and the envelope plasmid VSV-G. Transfections were performed for either six hours or overnight, after which the medium was replaced with virus collection medium (DMEM supplemented with 10% FBS and 1% bovine serum albumin; Cytiva HyClone).

Viral supernatants were collected 48–72 hours after medium replacement and centrifuged at 200 × g for 5 minutes to remove cellular debris. Clarified supernatants were aliquoted and stored at −80 °C until use.

Lentiviral titers were determined using a flow cytometry–based transduction assay in 384-well format. Briefly, 6,000 HEK293T cells were seeded per well in 384-well plates. The following day, approximately 10,000 cells were transduced by adding 0.5 µL of lentiviral supernatant to each well. Seventy-two hours post-transduction, culture medium was removed and cells were dissociated by adding 20 µL of 0.25% Trypsin-EDTA (Thermo Fisher Scientific) followed by incubation at 37 °C for 3 minutes. An additional 20 µL of FACS buffer (1× phosphate-buffered saline supplemented with 2% EDTA and 5% fetal bovine serum) was added, and cells were mixed thoroughly to ensure complete dissociation.

Cells were then analyzed by flow cytometry using the High Throughput Sampler (HTS) module of an LSRFortessa™ Cell Analyzer (BD Biosciences) at the University of Zurich Cytometry Core Facility. The proportion of transduced cells was quantified based on BFP fluorescence. Viral titers, expressed as transducing units per milliliter (TU/mL), were calculated using the formula TU = (P × N) / V, where P is the number of BFP-positive cells, N is the total number of cells at the time of infection, and V is the volume of viral supernatant added. Titers for the genome-wide screen were determined using HEK293T cells to enable high-throughput, plate-scale titer estimation, whereas titration for individual constructs used in downstream validation experiments was performed in U-251 MG cells to reflect the biological context of the screening system. To assess potential intra-plate gradients during viral production, titers were additionally visualized as a spatial heatmap and frequency distribution for a representative plate (Fig. 7A,B).

**Figure 7.**
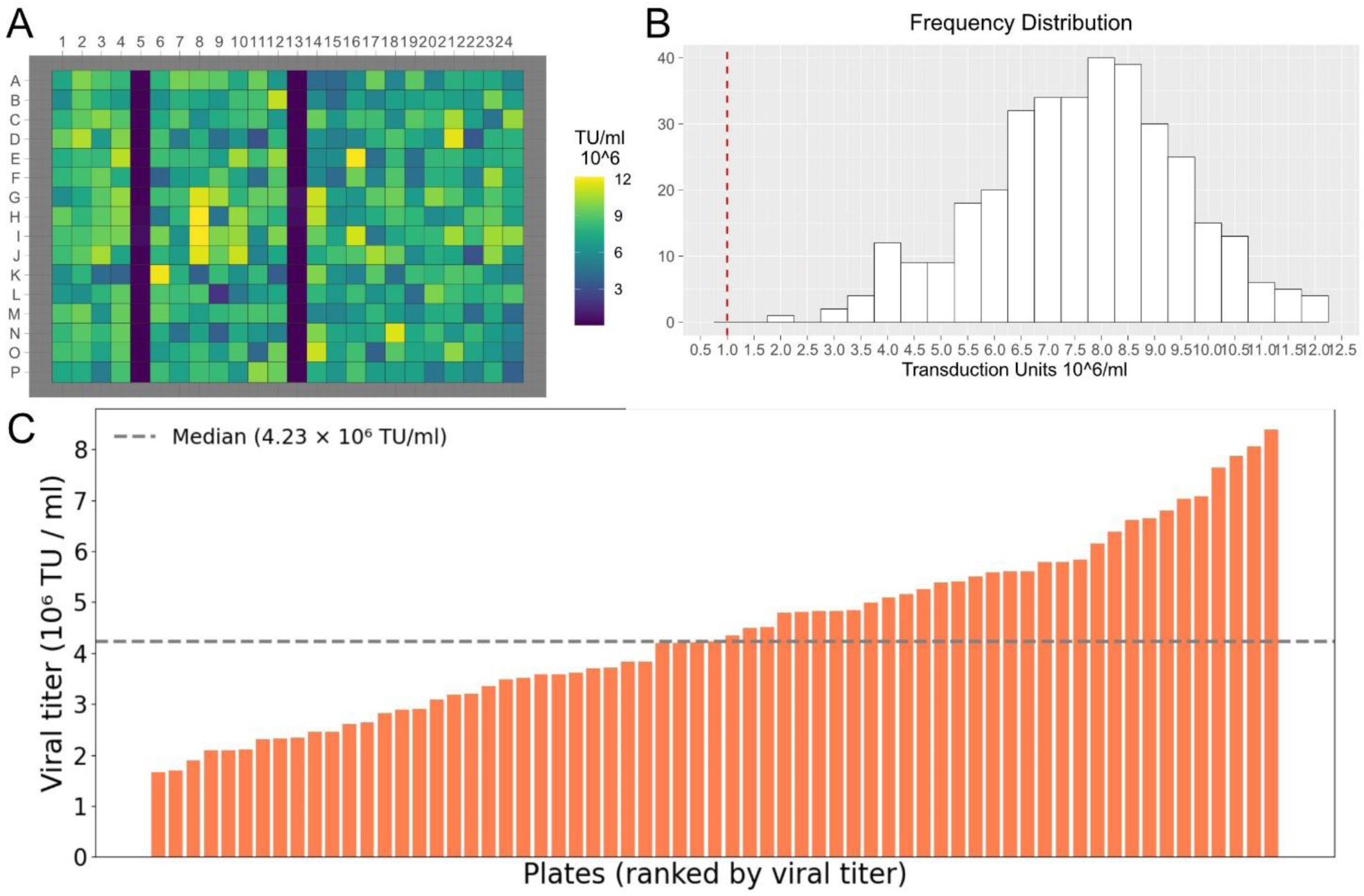
Plate-to-plate variability and intra-plate distribution of lentiviral titers. (A) Heatmap of lentiviral titers across a representative 384-well plate (plate 58). Columns 5 and 13 were reserved for control conditions (non-targeting and PRNP-targeting constructs) and contained cells and media but no CRISPR library transfection. Titers in these wells are therefore close to zero and are displayed accordingly in the heatmap. The remaining wells show measured viral titers across the plate, illustrating the spatial distribution of viral production. Although individual wells exhibit minor variability, the mean titer per plate (excluding control wells) was used to define the effective viral titer for that plate. (B) Frequency distribution of lentiviral titers from the representative plate shown in (A). The x-axis indicates viral titer (10⁶ TU/ml), and the y-axis indicates the number of wells per titer bin. (C) Mean lentiviral titers per production plate, excluding control wells, ranked from lowest to highest. Bars represent individual plates and are colored in golden orange. The dashed grey line indicates the median viral titer. Titers are expressed as 10⁶ transducing units (TU) per ml. For downstream screening, the mean titer of each plate (excluding control wells) was used to calculate viral input volumes required to achieve a multiplicity of infection (MOI) of 3 for transduction.

Lentiviral titers exhibited substantial plate-to-plate variability (Fig. 7C), with a mean titer of 4.41 × 10⁶ TU/ml (median 4.23 × 10⁶ TU/ml; SD 1.71 × 10⁶ TU/ml; CV 38.7%; range 1.66–8.38 × 10⁶ TU/ml), supporting the use of plate-level normalization strategies in downstream screening analyses.

### Genome wide CRISPRa screen

U-251 MG cells stably expressing dCas9-VPR were seeded into 384-well plates (PerkinElmer) at a density of 5,000 cells per well in a total volume of 30 µL using a MultiFlo FX Multimode Dispenser (Agilent). Plates were centrifuged at 10 × g for 30 s (Eppendorf 5804R) and incubated overnight in a rotating tower incubator (LiCONiC StoreX STX). After 18–20 h, cells were transduced with qgRNA-encoding lentiviruses at a multiplicity of infection (MOI) of 3 using an electronic multichannel pipetting system (ViaFlo, Integra). Each lentiviral construct encoded a quadruple-guide RNA cassette targeting a single human gene^14^. Continuous gentle rotation was used to promote uniform distribution of nutrients, oxygen, and lentiviral particles across the plate and to minimize evaporation-and temperature-related gradients across the plate. This approach reduces well position–dependent variability in cell growth and transduction efficiency.

To control for positional effects inherent to high-density plate formats, each 384-well plate incorporated a predefined layout strategy (Fig. 3A,B). Outer wells (columns 1 and 24) were not used for genetic perturbations; instead, they contained medium-only wells without cells and were reserved for TR-FRET normalization and background correction.

Genetic controls were placed in two internal columns (columns 5 and 13) to allow monitoring of spatial effects across the plate. Each of these columns contained 14 non-targeting controls (NTCs) and 14 PRNP-targeting positive controls arranged in alternating rows. The orientation of NTC and PRNP controls was mirrored between columns 5 and 13 (i.e., rows containing NTCs in column 5 contained PRNP-targeting controls in column 13, and vice versa). This mirrored arrangement enabled detection of row-and column-specific biases in the TR-FRET readout and allowed internal correction of spatial trends across the plate surface.

All remaining wells (excluding columns 1, 5, 13, and 24) were used for individual qgRNA perturbations targeting single genes. Each perturbation was tested in duplicate on independent plates rather than within the same plate. Well positions were preserved across replicate plates to control for spatial effects while allowing independent viral transduction and assay processing. This design ensured that replicate measurements captured biological and technical variability across separate experimental runs rather than only intra-plate variation.

Together, this plate configuration enabled (i) internal monitoring of assay performance using spatially distributed positive and negative controls, (ii) correction for edge and positional artifacts, and (iii) robust estimation of reproducibility across independent replicate plates. After incubation, culture medium was removed by plate inversion, and cells were lysed in 10 µL of lysis buffer containing 50 mM Tris–HCl (pH 8), 150 mM NaCl, 0.5% sodium deoxycholate, 0.5% Triton X-100, and EDTA-free cOmplete protease inhibitor (Roche). Plates were incubated on a ThermoMixer Comfort (Eppendorf) for 10 min at 4 °C with shaking at 450 rpm, followed by centrifugation at 1,000 × g for 1 min. Lysates were incubated at 4 °C for 2 h and centrifuged again under the same conditions.

PrP^C^ abundance was quantified using a solution-based TR-FRET immunoassay^16^. Five microliters of each FRET antibody pair were added to the clarified lysates to achieve final concentrations of 2.5 nM donor and 5 nM acceptor antibodies in 1× Lance buffer (PerkinElmer). The assay employed two anti-PrP antibodies recognizing distinct PrP^C^ epitopes^17^: POM2 conjugated to europium (Eu) as the donor fluorophore and POM1 conjugated to allophycocyanin (APC) as the acceptor. Eu–POM2 conjugation was performed as previously described^16^, and APC–POM1 coupling was carried out using the Lightning-Link APC labeling kit according to the manufacturer’s instructions.

For TR-FRET assay calibration and background correction, donor-only and acceptor-only antibody controls were included on each plate (Fig. 3B). Wells containing Eu–POM2 alone or APC–POM1 alone were used to quantify donor and acceptor background fluorescence and assess bleed-through and cross-excitation. Additional wells lacking both antibodies, containing only 1x Lance buffer, were included to measure baseline signal, while wells containing both antibodies served as positive FRET controls. This design enabled accurate background subtraction and calculation of net TR-FRET signal for PrP^C^ quantification across the plate. A representative plate-level heatmap of raw net TR-FRET signal, arranged according to the assay layout, is shown to illustrate the spatial distribution of the primary readout and the placement of internal controls used for quality metrics (Fig. 3C).

Plates were centrifuged and incubated overnight at 4 °C prior to signal acquisition. TR-FRET signals were measured using an EnVision multimode plate reader (PerkinElmer). The Eu donor was excited at 337 nm, and emission was recorded at 615 nm (donor) and 665 nm (acceptor)^16^.

### Computational processing and analysis of the genome-wide CRISPRa screen

Raw TR-FRET measurements from the genome-wide arrayed CRISPR activation (CRISPRa) screen were processed using a fully scripted Python analysis pipeline (Fig. 8) that converts plate-reader exports into per-perturbation effect sizes, statistical significance estimates, hit tables, and publication-quality figures. The pipeline operates on three primary data inputs: (i) raw TR-FRET plate exports organized as 384-well matrices, (ii) a layout/annotation table specifying the mapping between each plate–well position and its biological identity (gene target, plasmid ID, control status), and (iii) an Excel workbook containing genomic coordinates used for chromosomal localization visualizations. All processing steps are deterministic, version-controlled, and generate auditable intermediate files, enabling complete regeneration of derived results from raw exports.

**Figure 8.**
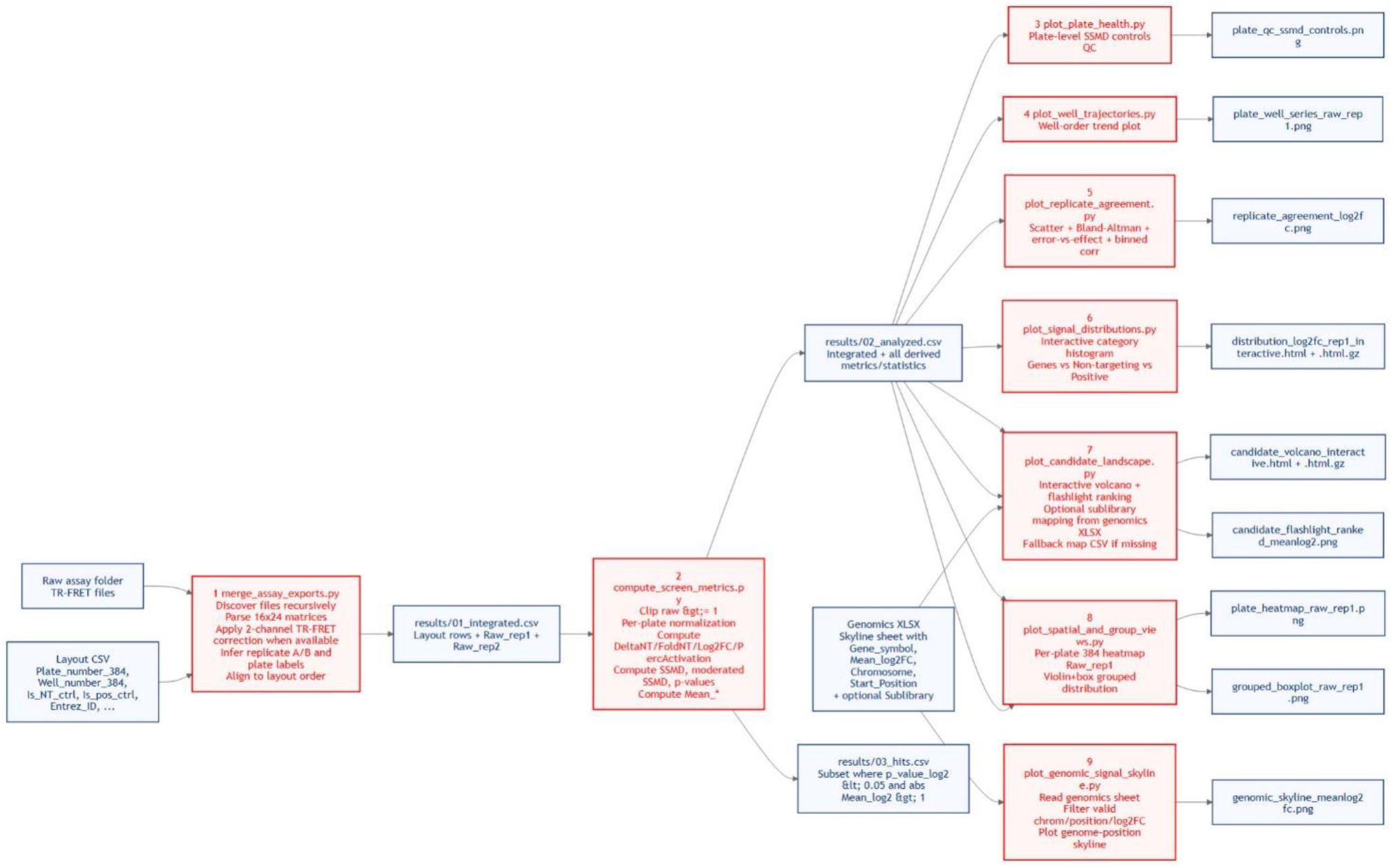
Computational workflow for processing and analysis of the genome-wide CRISPRa PrP^C^ screen. Schematic overview of the fully scripted Python pipeline used to process raw TR-FRET measurements and generate all primary screening results and figures. Raw assay exports from the genome-wide arrayed CRISPR activation screen are integrated with a layout/annotation CSV defining plate–well–gene mappings and control identities, and with an external genomics workbook providing chromosomal coordinates for genomic localization analyses. In Stage 1, raw TR-FRET plate files are recursively discovered, parsed as 384-well matrices, corrected using a two-channel TR-FRET model when available, and aligned to the layout scaffold to generate a unified per-well dataset (results/01_integrated.csv). In Stage 2, raw signals are clipped, normalized on a per-plate basis, and converted into derived metrics including log₂ fold-changes, percent activation, replicate means, and moderated SSMD-based p-values, yielding the analyzed screen table (results/02_analyzed.csv) and a thresholded hit list (results/03_hits.csv). Stages 3–8 generate quality-control, reproducibility, distribution, spatial, and candidate-level visualizations directly from the analyzed table, including plate QC metrics, replicate concordance plots, histograms, volcano plots, ranked effect summaries, and plate heatmaps. Stage 9 integrates genomic coordinate information to produce genome-wide localization (skyline) plots of PrP^C^ modulators. Script nodes are shown in red and data files in blue. All steps are deterministic and enable full regeneration of manuscript figures from raw assay exports.

*Input data and annotation scaffold*. The plate-to-gene (and plate-to-control) assignment is defined exclusively by a layout CSV containing one row per assayed well. Required columns include Plate_number_384, Well_number_384, Is_NT_ctrl, Is_pos_ctrl, and Entrez_ID, with optional metadata such as Gene_symbol, Plasmid_ID, and transcript or transcription start site identifiers. This layout table serves as the authoritative annotation scaffold for downstream integration and prevents implicit inference of well identities from file order or plate geometry alone.

Raw TR-FRET files are expected to contain 384-well matrices (rows A–P, columns 1–24 or 01–24) after skipping instrument-specific header lines. When present, TR-FRET exports may include two stacked 16 × 24 matrices corresponding to donor and acceptor emission channels, enabling application of a two-channel background-corrected TR-FRET model; otherwise, the first 16 × 24 block is used as a single-channel readout.

In the first processing stage (merge_assay_exports.py), the pipeline recursively discovers raw TR-FRET files from a user-specified directory, parses plate matrices, and applies a two-channel TR-FRET correction when both donor and acceptor channels are available. Replicate identity is inferred from filename suffix conventions (e.g., “A” for replicate 1 and “B” for replicate 2), and plate identifiers are extracted from standardized or legacy naming patterns.

Corrected plate-level signals are aligned to the layout scaffold by matching Plate_number_384 and ordering wells according to Well_number_384 using a row-major convention (A01 = 1 through P24 = 384). This explicit alignment step prevents plate or well misassignment and ensures that each measurement is attached to the intended gene or control defined in the layout table. The output of this stage is results/01_integrated.csv, which contains the layout metadata together with Raw_rep1 and Raw_rep2 for each perturbation.

In the second stage (compute_screen_metrics.py), raw TR-FRET signals are converted into normalized effect-size measures suitable for cross-plate comparison. Prior to log transformation, raw replicate values are coerced to numeric and clipped to a lower bound (minimum value of 1) to avoid undefined logarithms.

Normalization is performed on a plate-by-plate basis using a median baseline computed from all gene-annotated perturbations on each plate. From this baseline, the pipeline derives multiple representations of signal, including absolute deviation from baseline (DeltaNT), fold-change (FoldNT), log-transformed raw signal (Raw_log2), and log2 fold-change (Log_2_FC). In addition, percent activation (PercActivation) is calculated using PRNP-targeting positive controls as a reference scale, providing an interpretable, control-anchored metric of assay dynamic range.

For each perturbation, replicate measurements are summarized using the mean of replicate-normalized values (e.g., Mean_log2, computed from Log_2_FC_rep1 and Log_2_FC_rep2). Statistical significance is assessed using a moderated strictly standardized mean difference (SSMD) framework appropriate for duplicate measurements.

A per-row variance estimate is derived from the squared difference between replicates, and this variance is stabilized by incorporating a plate-level variance term estimated from non-targeting control wells. The resulting moderated SSMD statistic is then converted to a two-sided p-value by interpreting it under a Student’s t-distribution with one degree of freedom (df = 1), reflecting the two-replicate experimental design. This approach improves robustness of significance estimates in the low-replicate regime while preserving sensitivity to large, reproducible effects. The complete derived dataset—including normalized metrics, SSMD statistics, and p-values—is written to results/02_analyzed.csv.

Candidate hits are defined using a combined effect-size and significance threshold applied to the replicate-averaged log2 fold-change. By default, perturbations are classified as hits if p_value_log2 < 0.05 and |Mean_log2| > 1.0. The resulting hit list is exported to results/03_hits.csv, while the full screened population remains available in results/02_analyzed.csv for downstream visualization and reanalysis. This separation ensures transparent hit selection and allows alternative thresholds to be explored without reprocessing raw data.

All primary screen figures are generated directly from results/02_analyzed.csv using dedicated Python plotting scripts. Plate-level assay quality is evaluated by comparing PRNP-targeting positive controls and non-targeting controls using Z′-factor and SSMD-based metrics (Fig. 5A), summarized in plate-level quality control plots. Spatial and positional effects are inspected using well-order trajectories and representative plate heatmaps. Replicate agreement is assessed using scatter-based concordance plots of log2 fold-change values (Fig. 5B).

The global distribution of perturbation effects is visualized using interactive histograms stratified by gene perturbations and control classes (Fig. 5C). Candidate landscapes are summarized using volcano plots (Fig. 4), and ranked “flashlight” plots derived from Mean_log2 and the corresponding p-value column. Volcano plot inclusion is controlled by annotation: gene-annotated wells and designated control wells are included, while unannotated, non-control wells are excluded to avoid plotting normalization or empty positions.

*Genomic localization analysis.* Genomic localization visualizations (Fig. 9B-C) were performed using plot_genomic_signal_skyline.py, which consumes an external genomics Excel workbook containing, at minimum, Gene_symbol, Mean_log_2_FC, Chromosome, and Start_Position. After filtering for valid genomic annotations, effect sizes are plotted across chromosomal coordinates to visualize genome-wide spatial patterns of PrP^C^ regulation.

**Figure 9.**
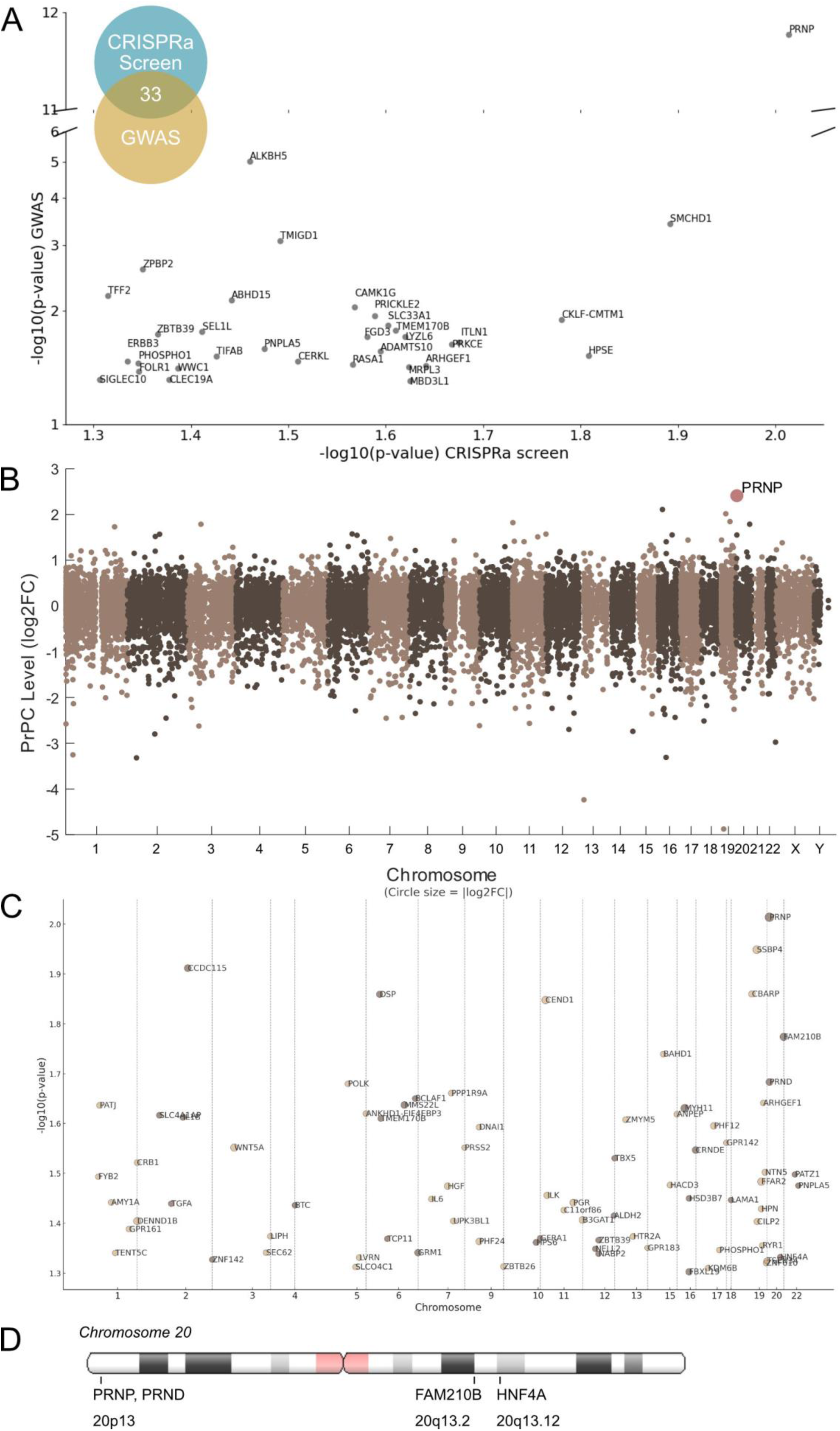
Genetic association and genomic localization of PrP^C^ modulators. (A) Overlap between CRISPRa hits and genome-wide association study (GWAS) loci for sporadic Creutzfeldt-Jakob disease (sCJD). Several PrP^C^ modulators map to regions associated with prion disease risk, suggesting functional convergence between physiological PrP^C^ regulation and genetic susceptibility. (B) Manhattan plot showing chromosomal position of the whole genome and their effect on PrP^C^ expression. No chromosome-specific trend was observed. (C) Mapping of the 80 PrP^C^ enhancers across chromosomes. Only four enhancers localize to chromosome 20, which contains the *PRNP* locus, suggesting a lack of enrichment. (D) Genomic positions of the four PrP^C^ enhancers on chromosome 20. Only PRND, which shares a bidirectional promoter with *PRNP*, may exert cis-regulatory effects. The others (FAM210B and HNF4A) are located distally.

*Interactive access and reanalysis.* To facilitate transparent reuse, interactive exploration, and reanalysis of the screening data, the full computational pipeline described here is also deployed as an online analysis platform at https://crispr4all.isab.science. This web-based interface implements the same data model, normalization procedures, statistical scoring, and visualization logic as the scripted Python pipeline used for the analyses in this study. Users can upload raw or processed screening datasets, apply standardized quality-control and normalization steps, reproduce key visualizations (including distribution plots, replicate concordance, volcano plots, and ranked candidate views), and export results for downstream analysis. The availability of both the fully scripted pipeline and the interactive platform ensures reproducibility while enabling broader community access to the dataset and analysis framework.

## Technical Validation

### CRISPRa functionality and assay optimization

To validate CRISPRa functionality in the screening system, U-251 MG cells stably expressing dCas9-VPR were transduced with quadruple-guide RNAs (qgRNAs) targeting the human prion protein gene *PRNP* or with non-targeting (NT) control qgRNAs. Western blot analysis demonstrated robust CRISPRa-mediated induction of PrP^C^ protein expression following *PRNP* activation (Fig. 2C), with maximal induction observed four days post-transduction. This time point was therefore selected as the standardized readout for all subsequent screening experiments.

Assay parameters were then optimized for large-scale, arrayed screening by systematically varying cell seeding density and lentiviral multiplicity of infection (MOI). Assay performance was evaluated using the Z′-factor, which quantifies the separation and dynamic range between positive and negative controls. Using *PRNP*-targeting qgRNAs as positive controls and NT qgRNAs as negative controls, a seeding density of 5,000 cells per well combined with an MOI of 3 consistently yielded high Z′-factor values (Fig. 2D) and strong separation between control populations. These optimized conditions were used for the entire genome-wide CRISPRa screen.

### Performance and reproducibility of the genome-wide CRISPRa screen

To ensure robust identification of genetic modulators of PrP^C^ abundance, multiple quality-control and validation measures were implemented throughout the arrayed genome-wide CRISPR activation (CRISPRa) screen. Each of the 22,442 CRISPRa perturbations was assayed in duplicate across independent 384-well plates, with well positions preserved between replicates to minimize spatial and batch effects. Non-targeting (NT) controls and *PRNP*-targeting positive controls were included on every plate to monitor assay performance and dynamic range.

Plate performance was monitored using 14 PRNP-targeting positive controls and 14 non-targeting controls per plate, positioned in columns 5 and 13. Assay quality was evaluated using the strictly standardized mean difference (SSMD) (Fig. 5A), calculated from *PRNP*-targeting and NT control wells on each plate. The majority of plates achieved Z′-factor values greater than 0.5, indicating excellent separation between positive and negative controls and suitability for high-throughput screening. Plate-to-plate reproducibility was assessed by calculating Pearson correlation coefficients (Fig. 5B) between replicate measurements, yielding a mean R² of 0.958 across the dataset, Visualization of TR-FRET measurements revealed two well-separated populations corresponding to *PRNP*-targeting and NT controls, while the majority of screened genes clustered near the plate median, consistent with most perturbations having no measurable effect on PrP^C^ abundance (Fig. 5C). During quality control analysis, we noted that the distribution of non-targeting control guides was shifted by 0.70355 log₂ units relative to the mean of all targeting guides. Importantly, this shift was consistent across replicates and did not affect the relative ranking of gene-level effects. To account for the systematic shifts in NT control distributions, PrP^C^ abundance values were normalized to the median signal of all genes on each plate. This normalization strategy reduces plate-specific bias and enables accurate estimation of relative effect sizes for individual genetic perturbations (Fig. 4).

The systematic shift observed for non-targeting controls likely reflects a technical rather than biological effect. Several plausible explanations can account for this phenomenon. First, the non-targeting controls may have originated from a different production or amplification batch than the bulk library, potentially introducing subtle differences in representation. Second, variations in storage conditions, freeze–thaw cycles, or plating density during library handling could have resulted in minor baseline abundance differences. Third, differences in cloning efficiency, titer normalization, or lentiviral preparation between control and targeting constructs could also contribute to a uniform offset. Notably, the shift was global and did not distort the variance structure or gene-level effect size estimates. Because hit calling was based on relative enrichment/depletion and replicated across independent screens, the offset does not materially affect the biological conclusions. Therefore, while the baseline displacement of non-targeting controls is visually apparent, it does not compromise the interpretability or reliability of the screening results.

### Genetic association analysis

To provide biological context for the dataset and to assess its relationship to known genetic risk factors for prion disease, we examined whether genes identified in the CRISPRa screen overlapped with loci associated with sporadic Creutzfeldt–Jakob disease (sCJD) in published genome-wide association studies (GWAS)^18^. Several CRISPRa hits mapped to genomic regions previously reported as sCJD risk loci (Fig. 9A), suggesting that a subset of genetic modulators of physiological PrP^C^ expression may intersect with pathways implicated in disease susceptibility.

To evaluate the overall correspondence between CRISPRa screen results and sCJD GWAS signals, we compared gene-wise statistical significance across datasets. Correlation analyses revealed no meaningful association between CRISPRa-derived effect sizes and GWAS p-values (Pearson r = – 0.00013, p = 0.9852; Spearman ρ = –0.0078, p = 0.2873), indicating that significance in one dataset does not predict significance in the other. Consistent with this observation, Kolmogorov–Smirnov testing showed no enrichment of GWAS-associated genes among CRISPRa hits, and Fisher’s exact tests performed across multiple significance thresholds did not identify significant overlap. A permutation test (p = 0.9852) and mutual information analysis (MI = 0.0077) further supported the statistical independence of the two datasets.

Together, these analyses indicate that, while individual overlaps between CRISPRa hits and sCJD-associated loci exist, there is no broad or systematic correspondence between regulators of PrP^C^ abundance identified in this screen and common genetic risk variants for prion disease. This distinction is important for appropriate interpretation and reuse of the dataset.

### Genomic localization analysis

To assess whether genes modulating PrP^C^ abundance exhibit non-random genomic distribution or chromosomal bias, all screened genes were mapped to their chromosomal locations together with their associated effects on PrP^C^ levels (Fig. 9B). This analysis revealed no evidence of chromosome-specific enrichment or clustering of PrP^C^ modulators, indicating that genetic regulation of PrP^C^ abundance is broadly distributed across the genome.

Because the CRISPRa system employs a strong transcriptional activator that can potentially influence genes located in close genomic proximity to the targeted locus, we specifically examined whether PrP^C^ upregulators identified in the screen were enriched near PRNP, the gene encoding PrP^C^, which is located on chromosome 20. Of the 80 PrP^C^ enhancers identified, only four mapped to chromosome 20 (Fig. 9C). Among these, only PRND (encoding the prion-like protein Doppel) is located adjacent to PRNP (Fig. 9D), making it a plausible candidate for local cis-regulatory effects.

In contrast, the remaining chromosome 20 enhancers, including FAM210B and HNF4A, are located approximately 50 Mb away from PRNP, rendering direct cis-regulatory interactions unlikely. These findings indicate that the vast majority of PrP^C^ enhancers identified in the CRISPRa screen do not act through genomic proximity to PRNP. Instead, they support a model in which PrP^C^ abundance is predominantly governed by trans-acting regulatory mechanisms, including transcriptional, signaling, and post-translational pathways.

### Validation of CRISPRa-mediated gene activation

CRISPRa functionality was validated by targeting the *PRNP* locus, which resulted in robust and reproducible induction of PrP^C^ protein levels as measured by both Western blotting and TR-FRET (Fig. 2C,D). Time-course experiments demonstrated that PrP^C^ expression increased progressively following transduction and reached a maximum four days post-transduction (Fig. 2C). This time point was therefore selected as the optimal readout window for the genome-wide CRISPRa screen.

The T.Gonfio CRISPRa library employs quadruple-guide RNA (qgRNA) constructs targeting transcription start sites and is designed to tolerate common human genetic polymorphisms. This architecture minimizes guide dropout, enhances activation potency, and promotes consistent gene induction across experimental conditions and cell lines. The high validation rate observed in downstream assays further supports the effectiveness and specificity of this CRISPRa-based gain-of-function strategy.

### Validation of screen hits

To independently validate the primary screening results, a subset of 50 candidate genes (22 PrP^C^ enhancers and 28 PrP^C^ reducers) was selected for validation testing. Candidates were chosen based on effect size in the primary screen and documented neuronal expression to ensure biological relevance.

PrP^C^ abundance was re-measured using two independent assays: secondary TR-FRET measurements and Western blotting. Secondary TR-FRET assays confirmed both the effect of PrP^C^ modulation for 45 of the 50 candidates (90%) (Fig. 6A). Effect sizes from the primary and secondary TR-FRET screens showed strong concordance (Fig. 6B, Pearson r=0.95), demonstrating high reproducibility of the primary genome-wide CRISPRa screen.

Western blotting provided antibody-based validation independent of the TR-FRET detection system and confirmed PrP^C^ upregulation for representative enhancer hits (Fig. 6C,D). PrP^C^ protein levels were detected using the POM2 antibody, with vinculin serving as a loading control. Western blot quantification was concordant with TR-FRET measurements (Fig. 6E), further supporting the reliability of the screening data.

Together, these validation approaches establish a high validation rate and demonstrate that the majority of curated hits represent bona fide genetic modulators of PrP^C^ abundance.

### Biological insight from GPCR-focused analysis of the CRISPRa screen

To further assess the biological coherence and interpretability of the genome-wide CRISPRa screen, we performed a complementary post hoc analysis focused on predefined functional gene classes. We specifically examined the G protein–coupled receptor (GPCR) sublibrary because GPCRs represent a well-annotated, pharmacologically tractable class of cell-surface signaling proteins with established roles in neuronal signaling, synaptic plasticity, and neurodegeneration, processes in which PrP^C^ has been repeatedly implicated.

Using a limma-based moderated t-statistic framework, GRM1, encoding metabotropic glutamate receptor 1 (mGluR1), emerged as one of the strongest and most reproducible PrP^C^ upregulators within the GPCR sublibrary (Fig. 10). The emergence of GRM1 is notable in light of prior studies demonstrating functional and physical interactions between PrP^C^ and glutamatergic signaling components, including metabotropic glutamate receptors, in neuronal physiology and excitotoxicity^19–21^. However, previous work has primarily positioned PrP^C^ as a modulator or signaling cofactor downstream of glutamate receptor activity^19–21^. In contrast, the present CRISPRa-based finding indicates that activation of GRM1 is sufficient to robustly increase PrP^C^ abundance, revealing a previously unrecognized upstream regulatory relationship.

**Figure 10.**
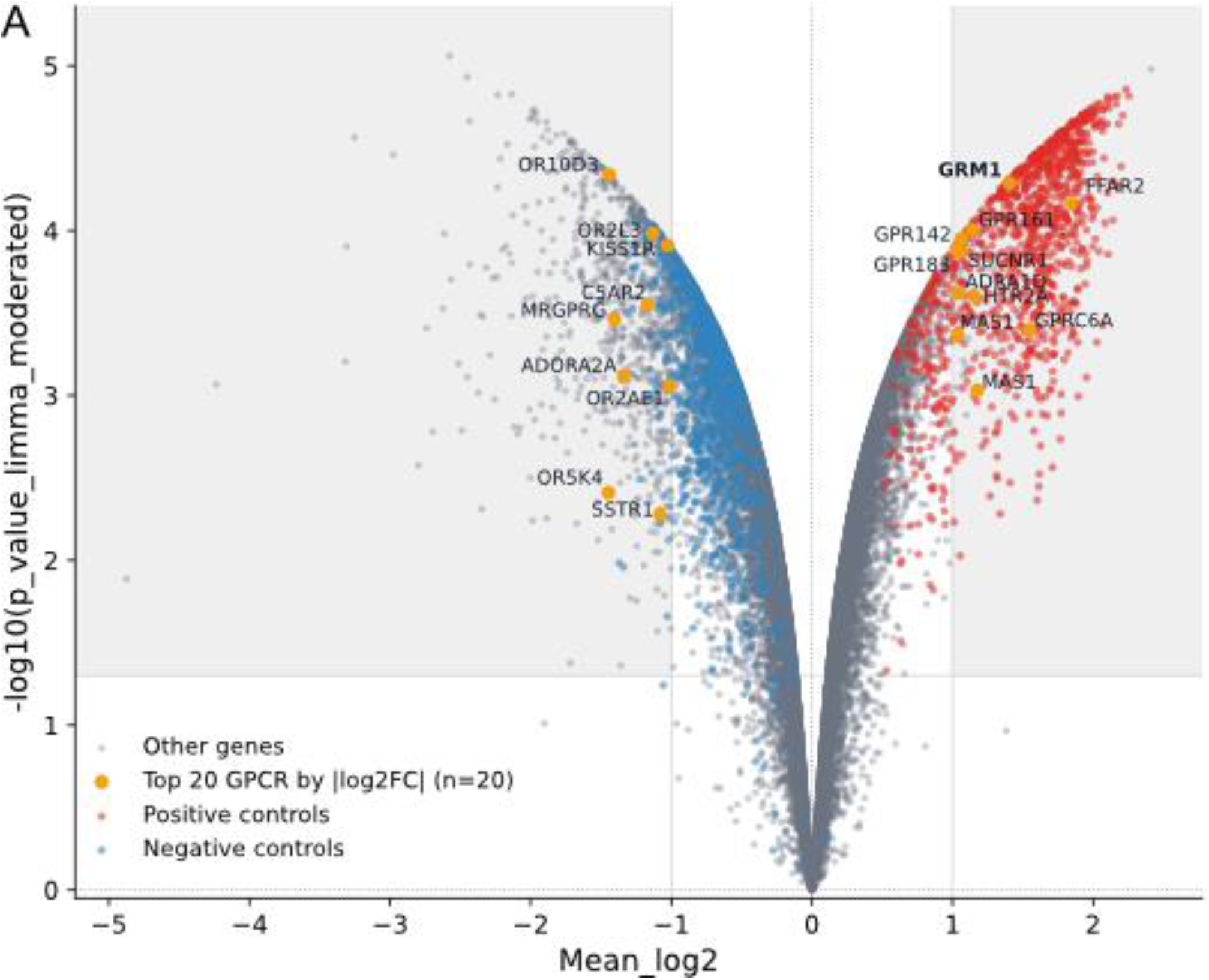
GPCR-focused reanalysis identifies GRM1 as a strong regulator of PrPᶜ abundance. (A) Volcano plot showing limma-based moderated t-statistics for genes within the GPCR sublibrary. The x-axis indicates log₂ fold-change in PrPᶜ abundance, and the y-axis shows –log₁₀ adjusted p-values derived from the limma model. The top 20 GPCR genes ranked by absolute log₂ fold-change are highlighted in yellow and labeled. GRM1 (metabotropic glutamate receptor 1) emerges as one of the strongest and most reproducible PrPᶜ upregulators within this functional gene class. Non-targeting (NT) control qgRNAs are highlighted in blue, and PRNP-targeting qgRNAs are highlighted in red.

Importantly, GRM1 was identified without targeted hypothesis testing and outside the primary hit-calling framework, underscoring both the depth of the dataset and the utility of orthogonal analytical approaches for extracting additional biological insight. Together, this result highlights glutamatergic signaling as a potential regulatory axis controlling PrP^C^ homeostasis and illustrates how reanalysis of the same screen with complementary statistical models can uncover distinct, biologically meaningful relationships.

### Data Records

All raw and processed datasets are provided in standardized formats with detailed annotations. This enables independent reanalysis, benchmarking, and integration with external datasets. Together, the extensive technical validation steps and high concordance across assays establish the reliability and utility of this dataset as a resource for the study of PrP^C^ homeostasis and intracellular protein regulation.

All data supporting the findings of this study are provided as raw inputs, processed tables, and supplementary source files to ensure full transparency and reproducibility. Raw TR-FRET plate-reader exports from the genome-wide CRISPRa screen, which serve as the primary input for generating the screen-level analyses and Figures 4–5, are available in the folder ScreenResults/PrPScreen_raw-data/TR-FRET. The corresponding plate and well annotation files defining gene identities, control positions, and plate layouts for all screened plates are provided in ScreenResults/PrPScreen_raw-data/LAYOUT. Processed outputs from the Python analysis pipeline—including the integrated per-well dataset, the fully analyzed dataset with normalized effect sizes and statistics, and the thresholded hit list—are provided as three Excel files generated directly by the pipeline. Supplementary datasets used for non-screen figures are organized according to figure order: TRFRET-Validation-RawData.xlsx contains secondary TR-FRET validation data used for Figures 6A–B; Titer-CRISPRaLibrary.xlsx contains lentiviral titer measurements used for Figure 7; PrP^C^_Hits_GWAS.xlsx provides data for GWAS overlap analyses shown in Figure 9A; and GeneticLocation.xlsx supplies genomic coordinate annotations used for chromosomal localization plots in Figures 9B–D. Uncropped Western blot source files used to assemble Figures 2 and 6 are provided as WB_Validation.afdesign and additional materials within the Optimization folder. Together, these datasets enable independent reanalysis, figure regeneration, and reuse of the data for future studies of PrP^C^ regulation and related cellular pathways

## Code Availability

All scripts used to process the raw TR-FRET measurements, perform plate-wise normalization, compute log₂ fold-changes and p-values, and generate the figures in this manuscript are available at: https://github.com/isab-science/Arrayed-CRISPR-screen-analyses-T.gonfio-T.spiezzo-.git. A schematic overview of the workflow is provided in Fig. 8.

In addition, the analysis pipeline described in the Methods is implemented in the CRISPR4ALL interactive platform (https://crispr4all.isab.science), which enables users to reproduce the analysis workflow, explore results, and perform custom analyses on the released datasets through a web interface.

## Contributorship statement

C.T.: designed, performed, contributed to and coordinated the genetic screens and their analyses, contributed to manuscript writing. H.W.: performed bioinformatic analysis, data curation and visualization; contributed to secondary screen, genetic association analysis and genomic localization analysis; experimental design and manuscript writing. V.B.: performed viral titration for the library.

S.M.: performed genetic association analyses comparing CRISPRa screen results with sCJD GWAS datasets. J-A.Y.: designed and produced the T.gonfio CRISPR activation library. A.A.: project design and supervision; secured funding; manuscript writing and correction, developed the Python pipeline for screen analysis.

## Funding and acknowledgements

A.A. is supported by a Distinguished Scientist Award of the NOMIS Foundation, grants from the Michael J. Fox Foundation (grant ID MJFF-022156 and grant ID MJFF-024255), the Swiss National Science Foundation (SNSF grant ID 179040 and grant ID 207872, Sinergia grant ID 183563), swissuniversities (CRISPR4ALL), the Human Frontiers Science Program (grant ID RGP0001/2022), the Parkinson Schweiz Foundation, the Uniscientia Foundation and the Creutzfeldt-Jakob Disease Foundation. H.W. is supported by the UZH Postdoc Grant (grant ID FK-25-058). The Institute for the Science of the Aging Brain is supported by a philanthropic donation by Adriano Aguzzi and by the Cantonal Government of St. Gallen, Switzerland. We thank Federico Bartolucci, Lorenzo Maraio, Ilan Margalith and Ana Marques for their help with lentivirus production, Andrea Armani and Geoffrey Howe for discussions, Lukas Frick and Athena Economides for assistance in bioinformatic analysis of the primary screen. The funders of the study had no role in study design, data collection, data analysis, data interpretation and manuscript writing. The corresponding author had full access to all the data in the study and had final responsibility for the decision to submit for publication.

