## Supplementary material for "Genome-wide arrayed CRISPR activation screen for prion protein modulators": candidate_flashlight_ranked_meanlog2_interactive.html

Candidate ranking flashlight


Sublibrary

Label points

Strongest modulators: Top 10
Strongest modulators: Top 20
Strongest modulators: Top 50
Gene list (textbox)

Apply
Clear

Publication SVG

Low (1200x800)
Medium (1800x1200)
High (2400x1600)
Export SVG
