## Supplementary material for "Genome-wide arrayed CRISPR activation screen for prion protein modulators": candidate_volcano_interactive.html

Interactive Volcano Plot


Primary p-value column (p\_value\_log2)
 limma moderated p (empirical-Bayes moderated t)

Experimental genes
 Positive controls
 Negative controls

Gene sublibrary

Highlight genes

Apply
Clear

Auto-label top genes

Largest |effect| (balanced +/-)
Smallest p-value
Strongest combo |effect| \* -log10(p)

Top 10
Top 20
Top 50
Apply
Clear

Publication SVG

Low (1200x800)
Medium (1800x1200)
High (2400x1600)
Export SVG

**Gene info**
Click a point to open external gene resources.

NCBI Gene
UniProt

**Legend and math**

- **Y-axis mode: Primary p** uses the analyzed table column `p_value_log2`.
- **Y-axis mode: limma p** uses empirical-Bayes moderated t-statistics (variance shrinkage across genes, limma-style).
- **Top labels: Largest |effect| (balanced +/-)** picks genes with largest `|Mean_log2|`, forcing both negative and positive sides when possible.
- **Top labels: Smallest p-value** picks lowest p under the currently selected Y-axis model.
- **Top labels: Strongest combo** uses score `|Mean_log2| * -log10(p_active)`. This is a ranking score, not a z-value.
- **Gene sublibrary filter** limits experimental genes and gene-hit overlays to one predefined sublibrary from the genomics workbook; controls remain controlled by category toggles.
- **Manual highlight** labels only user-entered genes; auto-top and manual highlight are independent overlays.
- **Export SVG** saves the currently visible state (Y mode, category toggles, labels/highlights) as a publication-ready vector figure.
