## Supplementary material for "Genome-wide arrayed CRISPR activation screen for prion protein modulators": replicate_agreement_log2fc_interactive.html

Replicate Diagnostics (Log2FC)


Interpretation Legend: Replicate Diagnostics

**What this figure measures:** replicate reproducibility across the same perturbations. Each point is one perturbation (gene or control).

**Panel 1 - Replicate Scatter:** x = rep1, y = rep2, with identity line `y = x`. Points near the line indicate agreement. Pearson `r` summarizes linear agreement, while outliers show discordant wells/genes. Controls (NT/positive) are highlighted for sanity-checking assay behavior.

**Panel 2 - Bland-Altman:** x = mean effect `(rep1 + rep2)/2`, y = difference `(rep2 - rep1)`. Center line is bias `mean(rep2 - rep1)`. Dotted lines are limits of agreement: `bias +/- 1.96 * SD(rep2 - rep1)`. This tests systematic shift and spread of replicate disagreement.

**Panel 3 - Error vs Effect Magnitude:** x = `|mean effect|`, y = `|rep2 - rep1|`, with a median trend. This quantifies heteroscedasticity (whether disagreement grows at stronger effects).

**Panel 4 - Binned Correlation vs |Effect|:** points are correlation estimates `corr(rep1, rep2)` computed within bins of `|mean effect|`. This shows whether reproducibility is stable across weak-to-strong signal regimes.

**Practical quality targets (rules of thumb):** Pearson `r >= 0.70` acceptable, `>= 0.80` good, `>= 0.90` excellent (assay-dependent). Bland-Altman bias near 0 and most points within LoA is desirable. Error-vs-effect trend should be flat or gently rising; steep growth suggests noisy extremes. Binned correlations should stay positive and preferably high across bins.

**What indicates concern:** broad scatter away from identity line, strong non-zero bias, fan-shaped Bland-Altman structure, rapidly increasing `|delta|` with effect size, or bin correlations collapsing toward 0/negative in high-effect bins.

**Interpretation note:** thresholds depend on assay dynamic range, replicate count, and control design. Always interpret with control separation (NT vs positive), hit concordance, and known biology.

Publication SVG

Low (1200x800)
Medium (1800x1200)
High (2400x1600)
Export SVG
