## Supplementary figures and images for "Genome-wide arrayed CRISPR activation screen for prion protein modulators"

### actin_2.6sec.01.03.21.Tif

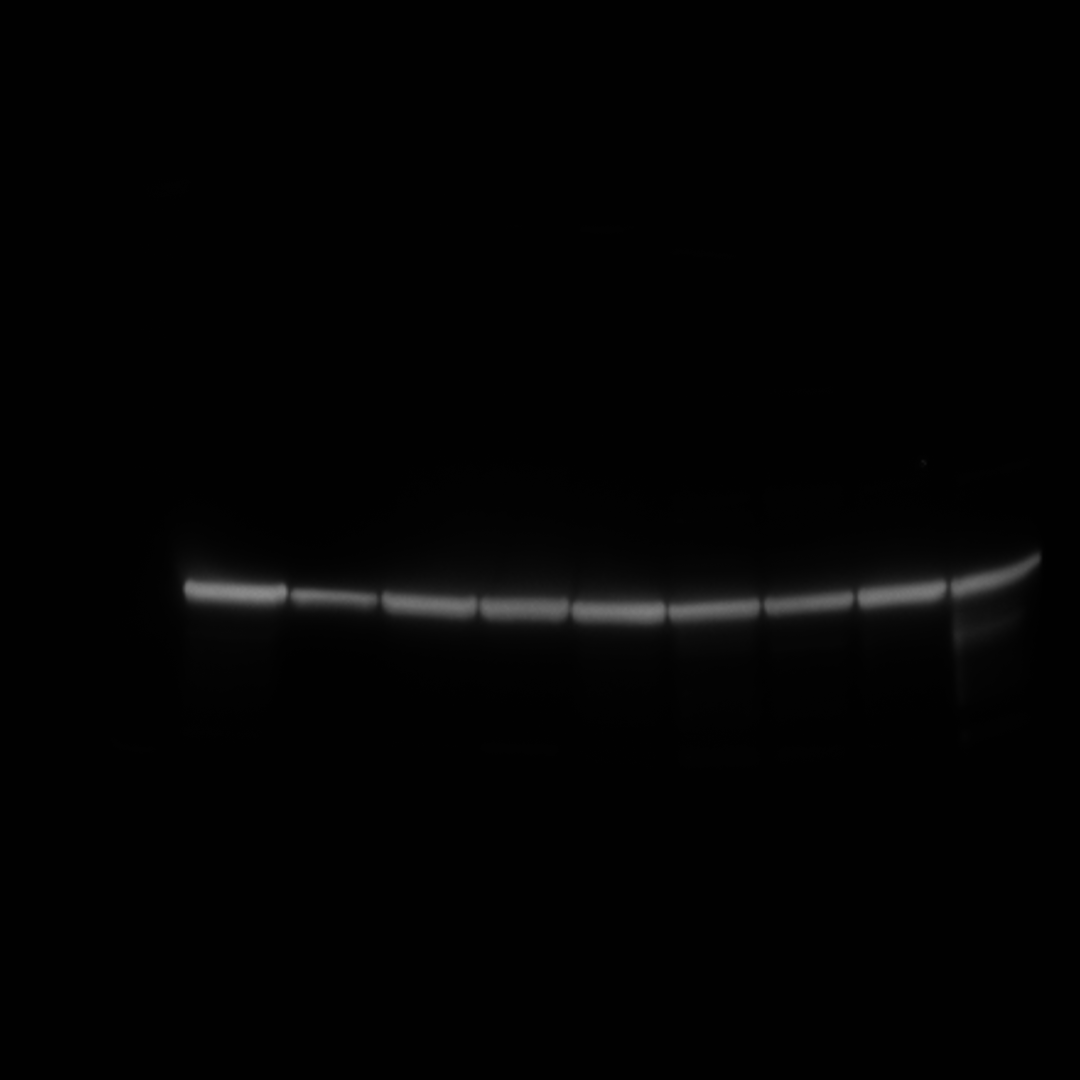

### ACTIN_U251MG_3-4-5-6d_notINF_NTa_2.30sec.TIF

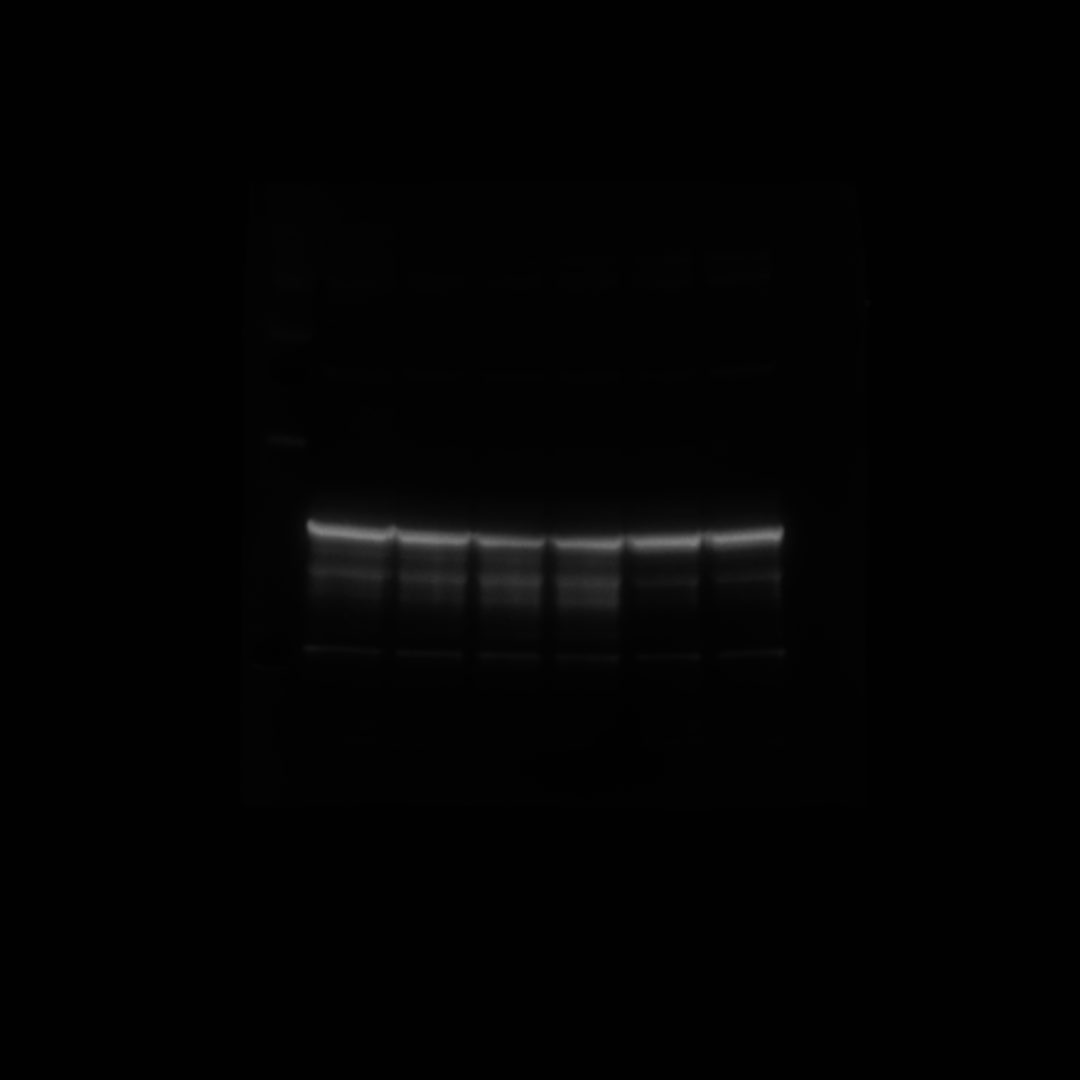

### candidate_flashlight_ranked_meanlog2.png

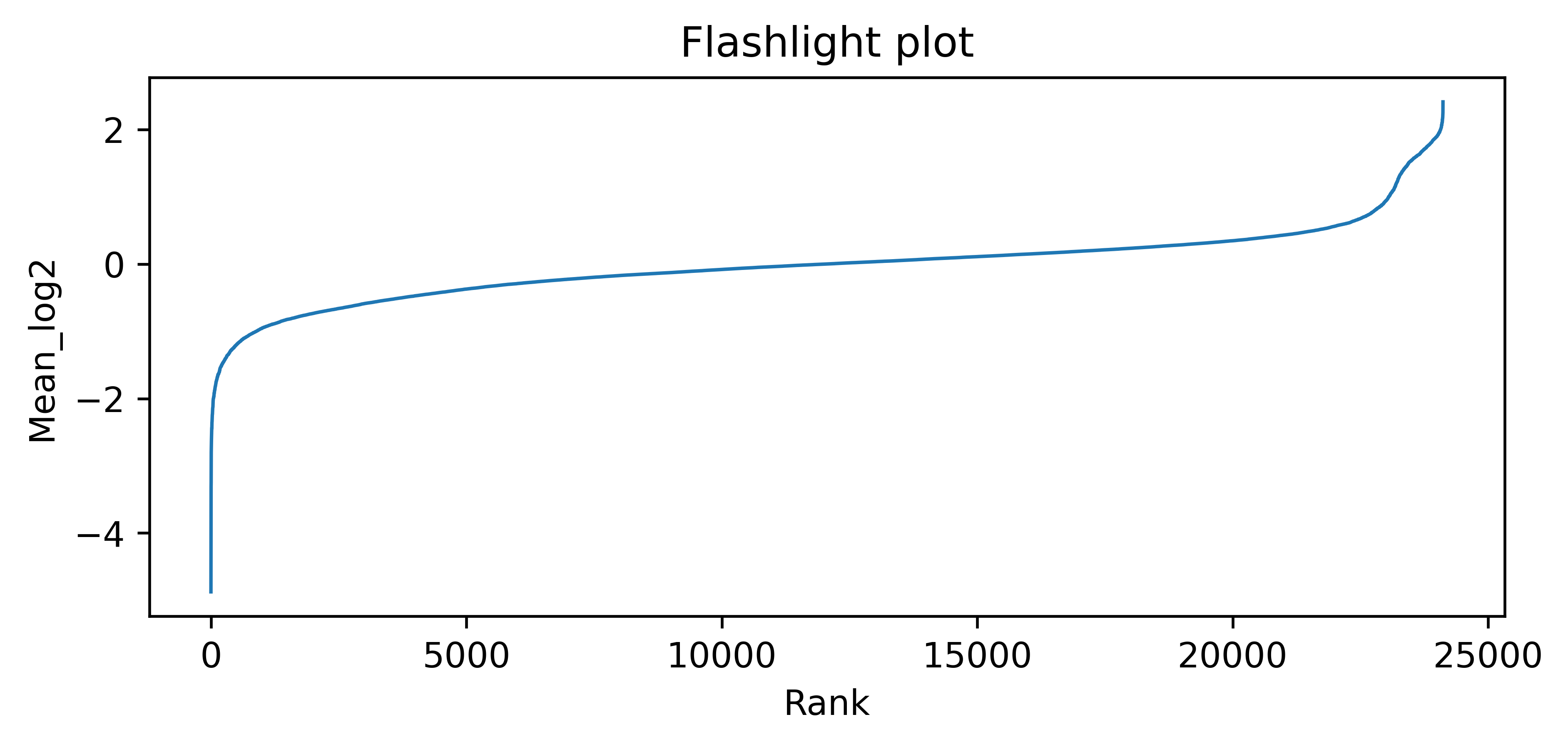

### grouped_boxplot_raw_rep1.png

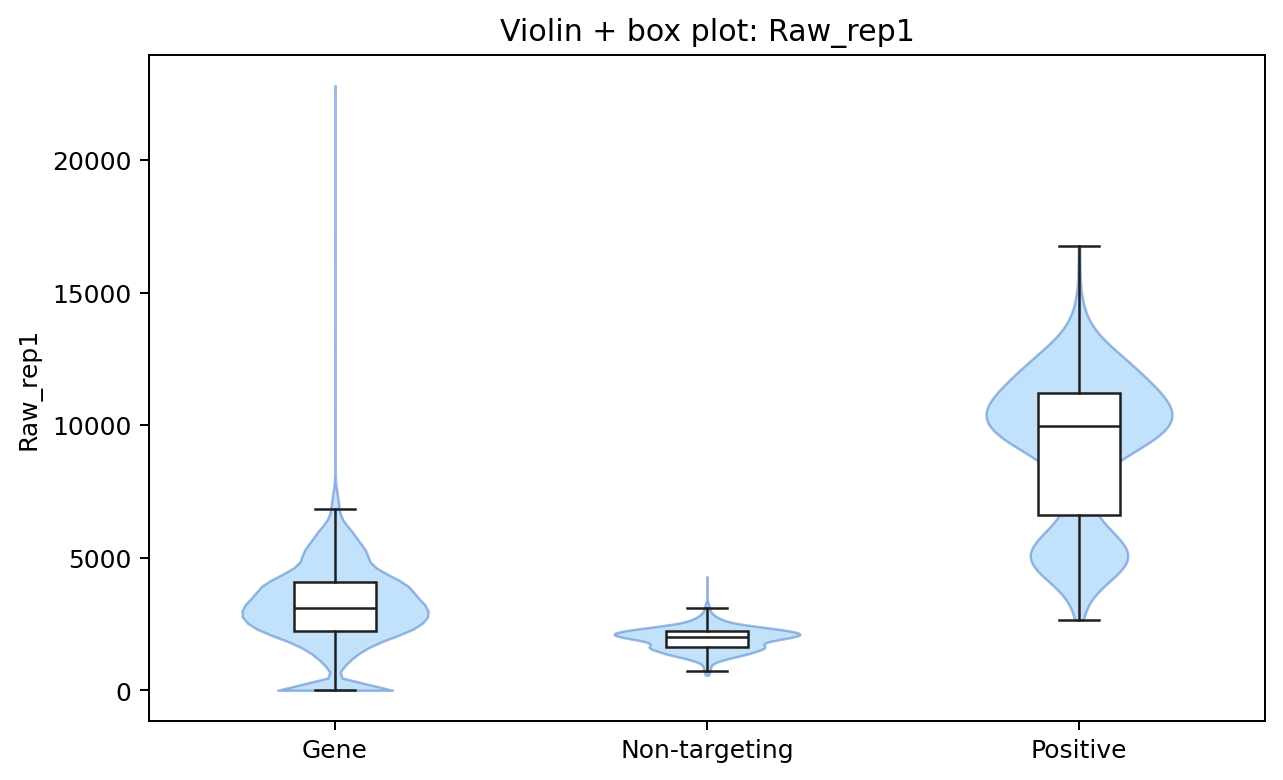

### plate_qc_ssmd_controls.png

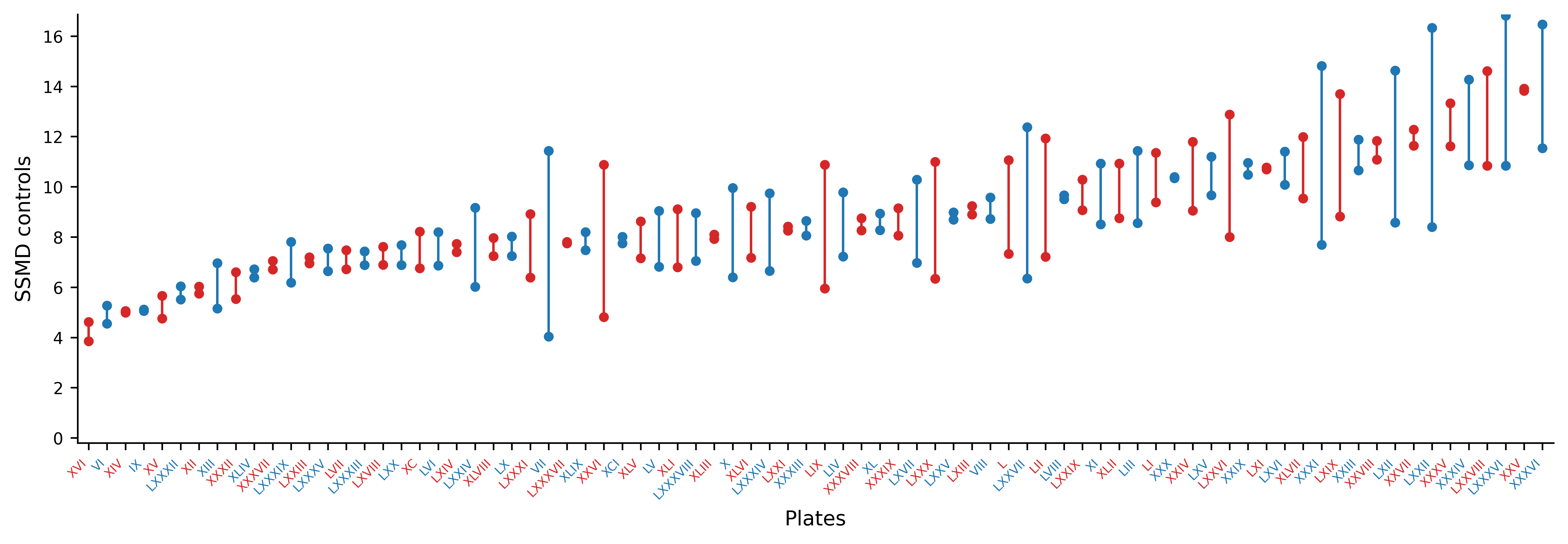

### plate_well_series_raw_rep1.png

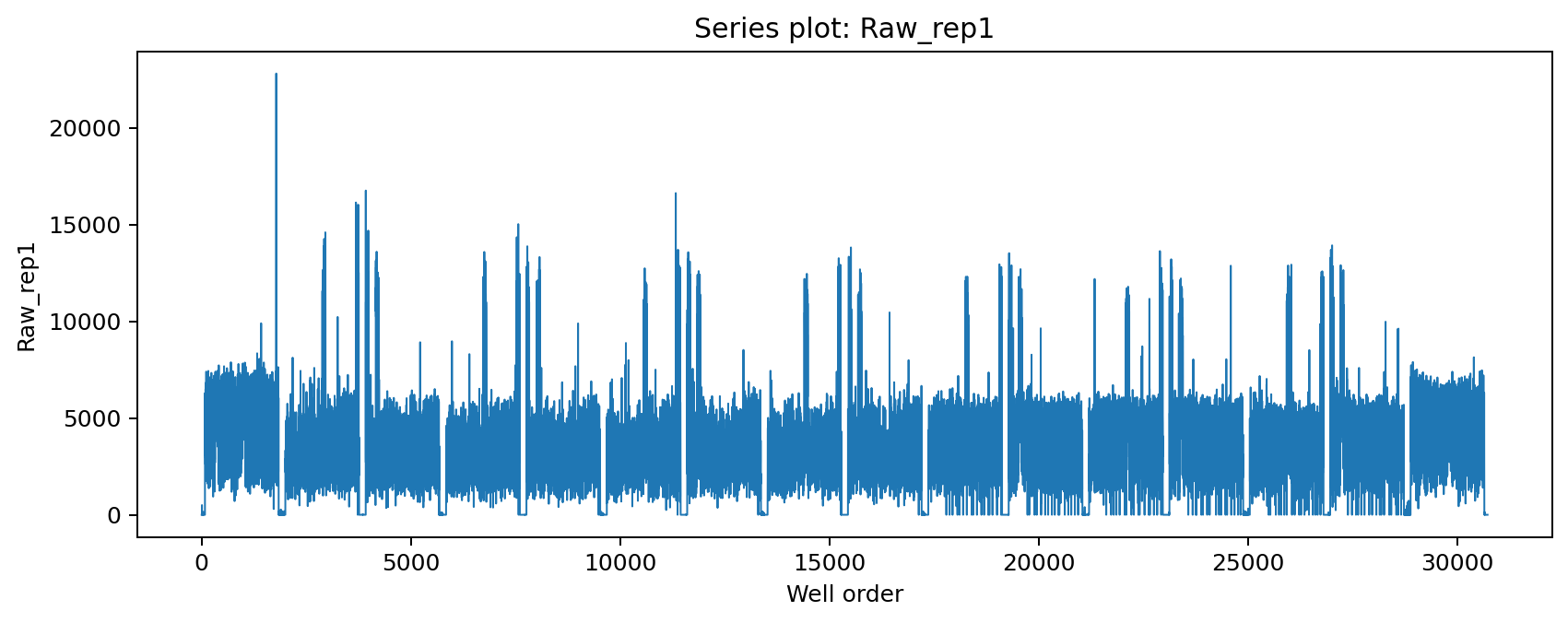

### POM2_5sec.Tif

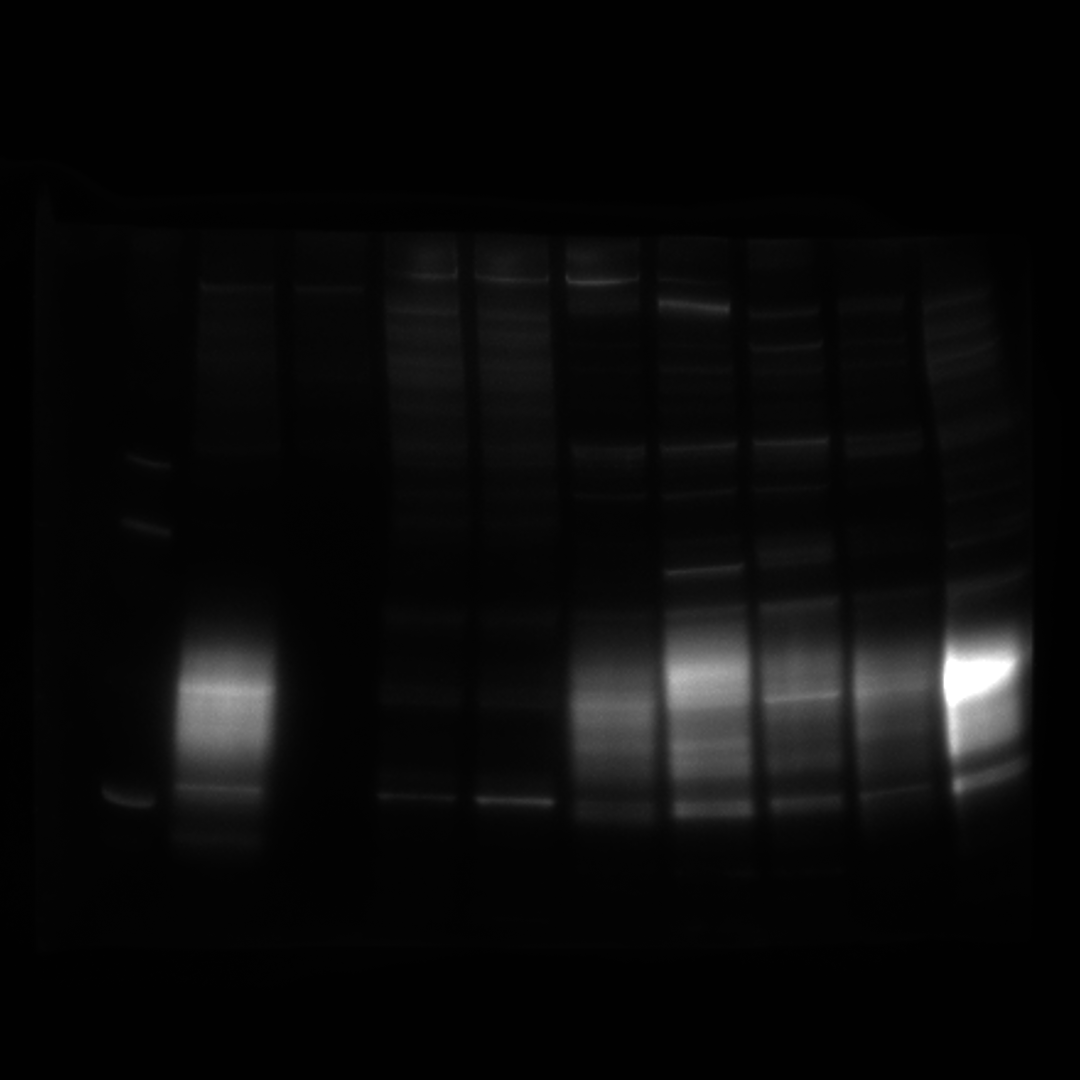

### replicate_agreement_log2fc.png

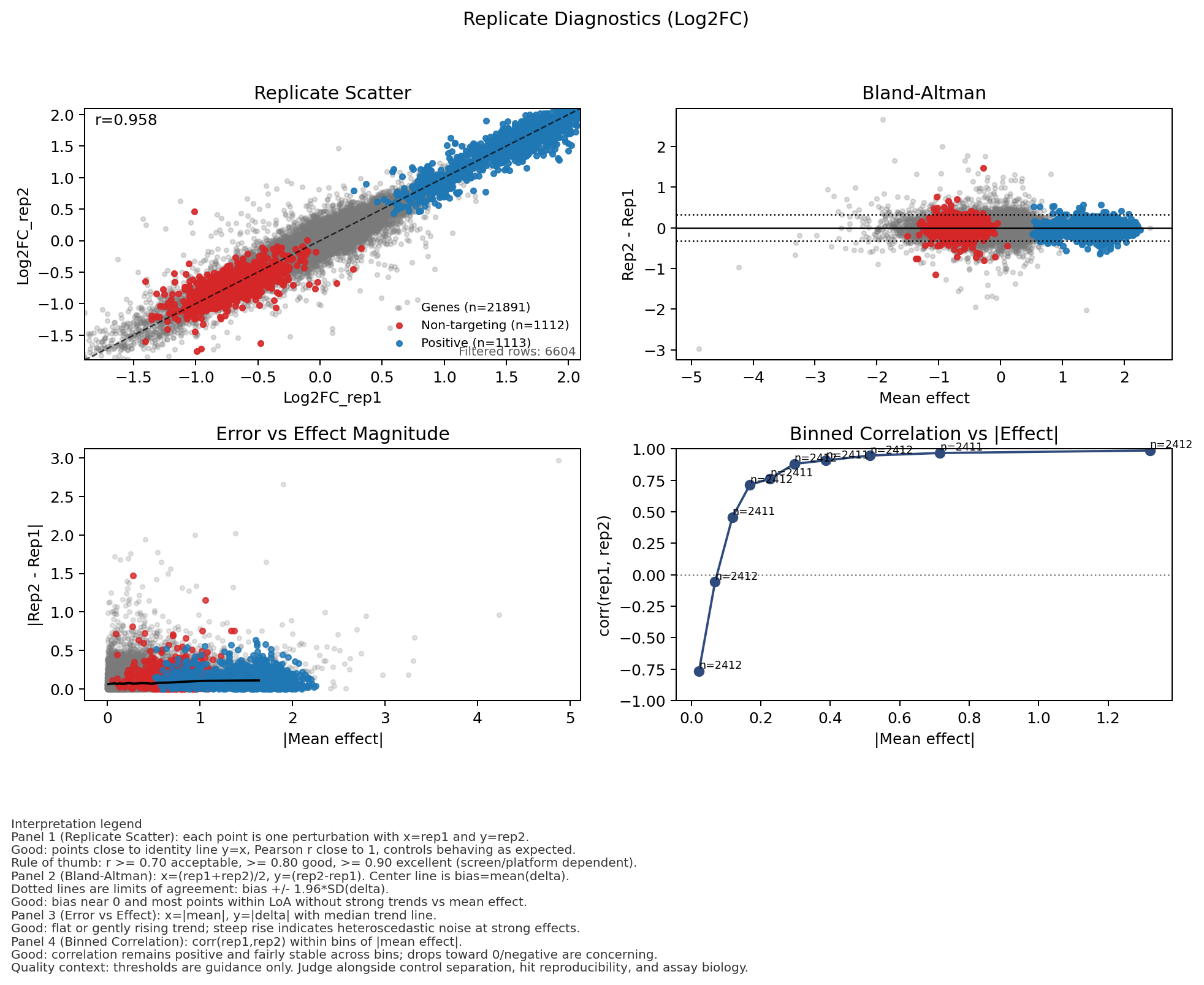

### U251MG_3-4-5-6d_nonInf_NTa_2Sum.TIF

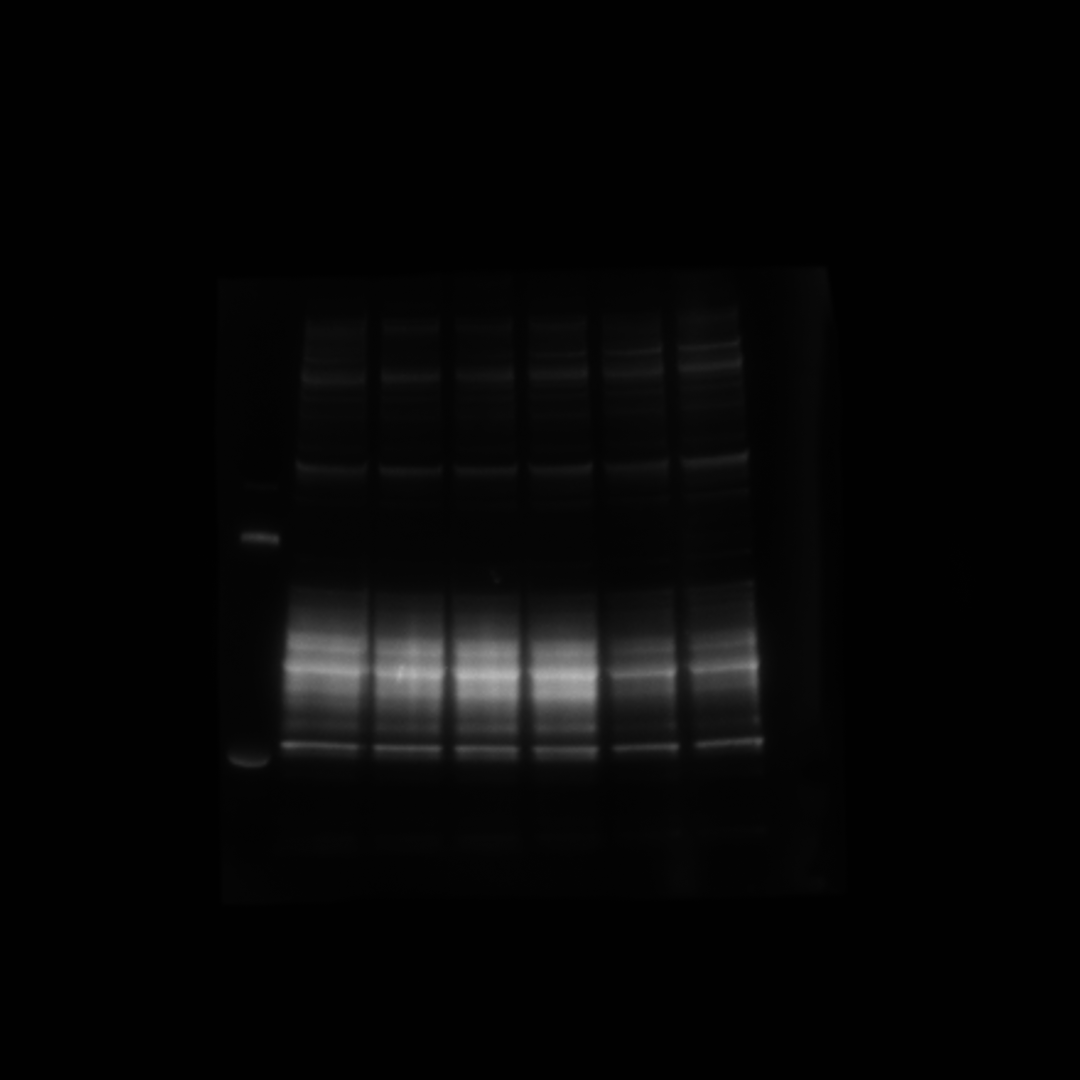
